# Polymerization cycle of actin homolog MreB from a Gram-positive bacterium

**DOI:** 10.1101/2022.10.19.512861

**Authors:** Wei Mao, Lars D. Renner, Charlène Cornilleau, Ines Li de la Sierra-Gallay, Sarah Benlamara, Yoan Ah-Seng, Herman Van Tilbeurgh, Sylvie Nessler, Aurélie Bertin, Arnaud Chastanet, Rut Carballido-López

## Abstract

In most rod-shaped bacteria, the actin homologue MreB is an essential component of the protein complex effecting cell wall elongation. The polymerization cycle and filament properties of eukaryotic actin have studied for decades and are well characterized. However, purification and *in vitro* work on MreB proteins have proven very difficult. Current knowledge of MreB biochemical and polymerization properties remains limited and is based on MreB proteins from Gram-negative species. In this study, we report the first observation of organized filaments and the first 3D-structure of MreB from a Gram-positive bacterium. We have purified MreB from the thermophilic *Geobacillus stearothermophilus* and shown that it forms straight pairs of protofilaments *in vitro*, and that polymerization depends on the presence of both lipids and nucleotide triphosphate. Two spatially close short hydrophobic sequences mediate membrane anchoring. Importantly, we demonstrate that unlike eukaryotic actin, nucleotide hydrolysis is a prerequisite for MreB interaction with the membrane, and that binding to lipids then triggers polymerization. Based on our results, we propose a molecular model for the mechanism of MreB polymerization.

## Introduction

Cytoskeletal proteins are known to polymerize into filaments that play critical roles in various aspects of cell physiology, including cell shape, mechanical strength and motion, cytokinesis, chromosome partitioning and intracellular transport. Prokaryotic cells contain homologs of the main eukaryotic cytoskeletal proteins, namely actin, tubulin and intermediate filaments (Cabeen & Jacobs-Wagner, 2010; Lin & Thanbichler, 2013; Shaevitz & Gitai, 2010), which were identified decades after their eukaryotic counterparts. In 2001, MreB proteins of the Gram-positive (G+) model bacterium *Bacillus subtilis* were found to form actin-like filamentous structures underneath the cytoplasmic membrane and to play a key role in the determination and maintenance of rod-shape (Carballido-Lopez, 2017; Jones *et al*, 2001). Soon after, the three-dimensional structure of one of the two MreB isoforms from the Gram-negative (G-) thermophilic bacterium *Thermotoga maritima* (MreB^Tm^) was solved (van den Ent *et al*, 2001), confirming its structural homology with actin (Bork *et al*, 1992). Besides, MreB^Tm^ in solution was shown to assemble into filaments similar to filamentous actin (F-actin) (van den Ent *et al*., 2001).

Research in the field of eukaryotic actin historically focused on elucidating structure-function relationships from *in vitro* studies. The availability of large amounts of soluble actin purified from several cell types since the 1940s enabled decades of mechanistic studies on actin polymerization (Pollard, 2016). In contrast, functional MreB from mesophilic bacteria proved particularly difficult to purify thwarting efforts to work with it *in vitro*. Instead, research on MreB primarily focused on cellular studies, driven by the advent of fluorescent microscopy in bacterial cell biology. Over the past two decades, the subcellular localization and dynamics of MreB have been described in several G- and G+ species (Billaudeau *et al*, 2017; Billaudeau *et al*, 2019; Dion *et al*, 2019; Harris *et al*, 2014; Hussain *et al*, 2018; Olshausen *et al*, 2013; Oswald *et al*, 2016; Ouzounov *et al*, 2016; Renner *et al*, 2013; Schirner *et al*, 2015). *In vivo*, MreB proteins form discrete membrane-associated polymeric assemblies along the cell cylinder that move processively around the rod circumference together with proteins of the cell wall (CW) elongation machinery (Domínguez-Escobar *et al*, 2011; Garner *et al*, 2011; van Teeffelen *et al*, 2011), forming the so-called Rod complex. The Rod complex motility is driven by CW synthesis (Domínguez-Escobar *et al*., 2011; Garner *et al*., 2011) and MreB assemblies self-align circumferentially, along their direction of motion (Billaudeau *et al*., 2019; Hussain *et al*., 2018). Recently, it was proposed that the specific intrinsic curvature of MreB polymers increases their affinity for the greatest concave (negative) membrane curvature within the cell (i.e. the inner surface of the rod circumference), accounting for their orientation (Hussain *et al*., 2018). The current model is that self-aligned MreB filaments restrict the diffusion of CW biosynthetic proteins in the membrane and orient their motion to insert new peptidoglycan strands in radial hoops perpendicular to the long axis of the cell, promoting the cylindrical expansion of rod-shaped cells (Domínguez-Escobar *et al*., 2011; Garner *et al*., 2011; Hussain *et al*., 2018). However, many questions remain to be answered. What prompts the assembly of MreB on the inner leaflet of the cytoplasmic membrane? What is the architecture of the membrane-associated MreB polymeric assemblies and how is it controlled? How is their distribution along the cell cylinder regulated? What is the length of individual MreB filaments within these assemblies and how is it controlled? Are the filaments stable? Do they exhibit turnover (treadmill) like actin filaments? *In vivo*, the length of MreB filamentous assemblies can be affected by the intracellular concentration of the protein (Billaudeau *et al*., 2019; Salje *et al*, 2011), but seems to have little impact on MreB function (Billaudeau *et al*., 2019). No turnover of MreB assemblies was detected *in vivo*, at least relative to their motion around the cell circumference (Domínguez-Escobar *et al*., 2011; van Teeffelen *et al*.,2011). Therefore, MreB polymers are believed to be quite stable despite their dynamic behavior in the cell. To elucidate in detail the molecular mechanisms underlying the functions of MreB, it remains necessary to understand their biochemical and polymerization properties. The majority of biochemical and structural studies on MreB proteins originally focused on the highly soluble G- MreB^Tm^ (Bean & Amann, 2008; Esue *et al*, 2005; Esue *et al*, 2006; Popp *et al*, 2010b; van den Ent *et al*., 2001; van den Ent *et al*, 2010). The tendency to aggregation upon purification hampered most *in vitro* studies of MreBs from other species (Dersch *et al*, 2020; Gaballah *et al*, 2011; Mayer & Amann, 2009). More recently, MreBs from several G- bacteria and from the wall-less bacterium *Spiroplasma citri* (MreB5^Sc^) could be purified in a functional soluble form, albeit in much lower quantities than MreB^Tm^ (Harne *et al*, 2020; Maeda *et al*, 2012; Nurse & Marians, 2013; Pande *et al*, 2022; Salje *et al*., 2011; van den Ent *et al*, 2014). Direct binding to the cell membrane was shown for MreB from the G- *Escherichia coli* and *T. maritima* (Salje *et al*., 2011). The N-terminal amphipathic helix of *E. coli* MreB (MreB^Ec^) was found to be necessary for membrane binding and also to cause the full-length purified protein to aggregate (Salje *et al*., 2011). Although this N-terminal amphipathic helix is dispensable for polymerization, it is required for proper function of MreB^Ec^ *in vivo* (Salje *et al*., 2011). MreB^Tm^ is devoid of such an N-terminal amphipathic helix, but instead possesses a small hydrophobic loop promoting membrane insertion that protrudes from the monomeric globular structure and was shown to also mediate membrane binding (Salje *et al*., 2011).

Altogether, *in vitro* work on MreBs from G- bacteria has shown that MreB polymerizes into straight double filaments in the presence of nucleotides, both in solution and on lipid membrane surfaces (Harne *et al*., 2020; Salje *et al*., 2011; van den Ent *et al*., 2014; van den Ent *et al*., 2010), and that filaments can assemble into larger sheets by lateral interactions (Esue *et al*., 2005; Esue *et al*., 2006; Harne *et al*., 2020; Nurse & Marians, 2013; Popp *et al*., 2010b; van den Ent *et al*., 2001; van den Ent *et al*., 2014). Furthermore, work on *Caulobacter crescentus* MreB (MreB^Cc^) and MreB^Ec^ indicated an antiparallel arrangement of the straight pairs of protofilaments (van den Ent *et al*., 2014), in sharp contrast to the helical parallel pairs of protofilaments (double helix) characteristic of F-actin (Pollard, 1990). While the parallel arrangement of a protofilament doublet generates polarity and allows the characteristic treadmilling of F-actin (Stoddard *et al*,2017), the antiparallel arrangement in MreB protofilaments suggests a bidirectional polymerization/depolymerization mechanism (van den Ent *et al*., 2014). The directionality and the kinetics of MreB polymerization, as well as the role of nucleotides in this process remain to be shown. ATPase activity has been reported in solution for MreB^Tm^, MreB^Ec^ and, more recently, for MreB5^Sc^ (Esue *et al*., 2005; Esue *et al*., 2006; Nurse & Marians, 2013; Pande *et al*., 2022; Popp *et al*., 2010b). However, the need for nucleotide binding and hydrolysis in polymerization remains unclear due to conflicting results, *in vivo* and *in vitro*, including the ability of MreB to polymerize or not in the presence of ADP or the non-hydrolysable ATP analogue AMP-PNP (adenylyl-imidodiphosphate). In addition, no electron microscopy (EM) images of protofilaments or atomic views of MreB from a G+ bacterium have been reported to date; all available EM and structural data are from G- species. In G+ bacteria, MreB proteins presumably have no N-terminal amphipathic helix (Salje *et al*., 2011), and the genome usually encodes several MreB isoforms (in contrast to G- that usually get by with a single *mreB* paralog), that may be related to their thicker and more complex CW structure (Chastanet & Carballido-Lopez, 2012). Inter- and intra-species differences in MreBs may exist at the structural or biochemical level, leading to differences in molecular interactions or biological functions.

In this study, we aimed to decipher for the first time fundamental structural and biochemical properties of MreB from a G+ bacterium. We successfully purified a soluble form of MreB from the G+ thermophilic bacterium *Geobacillus stearothermophilus* (MreB^Gs^) and elucidated its crystal structure, confirming the classical actin/MreB fold. Polymerization assays showed that MreB^Gs^ forms straight pairs of protofilaments in the presence of lipids and nucleotide triphosphate (either ATP or GTP). MreB^Gs^ does not polymerize in free solution like its G- counterparts. We have also shown that the interaction with lipids is mediated by two spatially close hydrophobic motifs in MreB^Gs^ monomers. Importantly, nucleotide hydrolysis was required for filament formation, in contrast to actin, which polymerizes spontaneously under physiological salt conditions and subsequently hydrolyzes ATP within the filament to promote depolymerization. Our results shed new light on the polymerization mechanism of MreB proteins.

## Results

### Crystal structure of *G. stearothermophilus* MreB

To overcome the notorious aggregation issues of MreB from mesophilic bacteria, we cloned and purified MreB from the thermophilic G+ bacterium *G. stearothermophilus* (MreB^Gs^). We chose *G. stearothermophilus* because of its proximity to the *Bacillus* genus and because of the highly conserved sequence of MreB^Gs^ compared to MreB from the model G+ bacterium *B. subtilis* (MreB^Bs^). MreB^Bs^ is more closely related to MreB^Gs^ (85.6 % identity and 92.6 % similarity) than to MreB of G- for which biochemical or structural data are available (either the thermophilic *T. maritima* with 55.8 % identity, or the mesophilic *C. crescentus*, 56.9 % identity and *E. coli*, 55.2 % identity) (Fig. S1).

MreB^Gs^ was purified to homogeneity following a two-step procedure (see Materials and Methods). The protein could be purified in a soluble form (Fig. S2A and C) and remained functional for polymerization at concentrations below 13 μM (0.5 mg/mL). When stored at higher concentrations or conserved at 4°C, MreB^Gs^ rapidly aggregated (Fig. S2A and B) and could not be recovered in a monomeric state, which is consistent with the known tendency of MreB proteins to aggregate.

The purified MreB^Gs^ protein was crystallized and the structure of its apo form was solved at 1.8 Å resolution (Protein Data Bank [PDB] ID 7ZPT). The crystals belong to the monoclinic P2_1_ space group and contain one molecule of MreB^Gs^ per asymmetric unit (Table S1). Monomers of apo MreB^Gs^ display the canonical fold of actin-like proteins, characterized by four subdomains IA, IIA, IB and IIB (Fig. 1A). One of the most similar structures to apo MreB^Gs^ is the apo form of MreB^Tm^ (PDB ID 1JCF, (van den Ent *et al*., 2001), with a rmsd of 1.92 Å over 305 superimposed Cα atoms and a Z-score of 16.0. Superimposition of the two proteins (Fig. 1A) revealed that MreB^Gs^ is in a slightly more open conformation than MreB^Tm^, mainly due to a movement of domain IB, which is the less conserved within the actin superfamily of proteins. Loop β6-α2, which connects subdomains IA and IB and closes the nucleotide-binding pocket is partially disordered in apo MreB^Gs^. In domain IA, the hydrophobic loop α2-β7, which has been shown to be involved in MreB^Tm^ membrane binding (Salje *et al*., 2011) and is 2 residues longer in MreB^Gs^ (Fig. S1), displays a distinct conformation, packed on the N-terminal extremity of the polypeptide chain.

**Figure 1.**
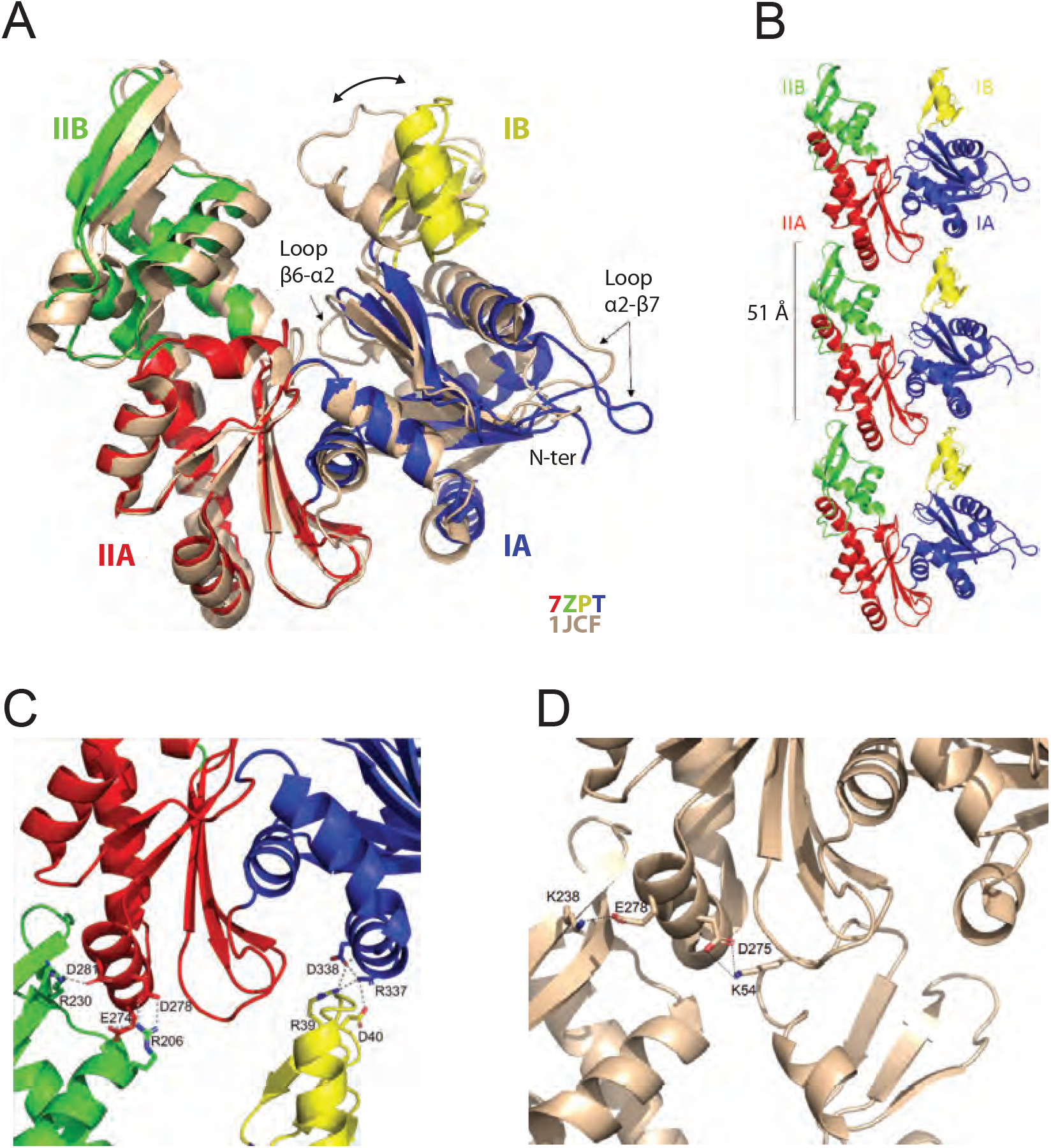
Crystal structure of the apo protofilament of MreB from *G. stearothermophilus*. (**A**) Crystal structure of apo MreB^Gs^ (PDB ID 7ZPT), colored by subdomains, superimposed on that of apo MreB^Tm^ (PDB ID 1JCF), in beige. The sequence similarity between the two proteins is 55.8%. Subdomain IA (blue) of MreB^Gs^ is formed by residues 1-32, 66-145 and 315-347; subdomain IB (yellow) by residues 33-65; IIA (red) by residues 146-181 and 246-314 and IIB (green) by residues 182-245. Superimposition of the two forms highlights the distinct positions of loops β6-α2 and α2-β7 as well as the movement of domain IB (two-headed arrow) resulting in slightly distinct subunit interaction modes as shown in panel C. (**B**) Protofilament structure of apo MreB^Gs^. Three subunits of the protofilament formed upon crystal packing are displayed as cartoon and colored by subdomains. The subunit repeat distance is indicated. (**C**) Close view of the MreB^Gs^ intra-protofilament interface. The two subunits are colored by subdomains as in panel A, and shown as cartoons. Residues involved in putative salt bridges (gray dashed lines) are displayed as sticks colored by atom type (N in blue and O in red) and labeled. (**D**) Close view of the MreB^Tm^ intra-protofilament interface (PDB ID 1JCF). The two subunits are colored in beige as in panel A, and shown as cartoons. Residues involved in putative salt bridges (gray dashed lines) are displayed as sticks colored by atom type (N in blue and O in red) and labeled.

Crystal packing analysis revealed that MreB^Gs^ molecules associate into straight protofilaments (Fig. 1B) characterized by a subunit repeat distance of 51 Å, similar to that observed in protofilaments of crystal structures of other actin homologs (Harne *et al*., 2020; Pande *et al*., 2022; Roeben *et al*, 2006; van den Ent *et al*., 2014). However, because of the open conformation of MreB^Gs^ (Fig. 1A), the interaction mode of the subunits observed in MreB^Gs^ protofilaments (Fig. 1C) is slightly different from that observed in protofilaments of MreB^Tm^ (Fig. 1D) (van den Ent *et al*., 2001), with domain IB interacting only with domain IA and not with domain IIA. While each interface in the MreB^Gs^ protofilament (Fig. 1C) is characterized by a solvation energy gain Δ^i^G of −7.1 kcal/mol, this value reaches −12.4 kcal/mol for MreB^Tm^ (PDB ID 1JCF) and −9.5 kcal/mol for MreB^Cc^ (PDB ID 4CZI), suggesting that the apo form of MreB^Gs^ forms less stable protofilaments than its G- homologs.

### MreB^Gs^ polymerizes into straight pairs of protofilaments in the presence of lipids

Next, we investigated the polymerization of MreB^Gs^ by EM of negatively stained samples. No filaments were observed under conditions in which MreB proteins from G- bacteria have been shown to polymerize in solution (Esue *et al*., 2005; Esue *et al*., 2006; Maeda *et al*., 2012; Nurse & Marians, 2013; Salje *et al*., 2011; van den Ent *et al*., 2001) (Fig. 2A and Table S2), suggesting that the purified protein was either nonfunctional for self-assembly or that a critical factor was missing. *In vivo*, MreB^Bs^ forms membrane-associated nanofilaments (Billaudeau *et al*., 2019; Hussain *et al*., 2018; Jones *et al*., 2001), and MreB filaments from G- bacteria have been shown to have intrinsic affinity for membranes (Garenne *et al*, 2020; Maeda *et al*., 2012; Salje *et al*., 2011; van den Ent *et al*., 2014). We hypothesized that the presence of lipids might be a prerequisite for the assembly of MreB^Gs^ polymers. On a lipid monolayer of total *E. coli* lipid extract, MreB^Gs^ readily formed filaments in the presence of ATP, which would not be observed without the biomimetic membrane (Fig. 2A). Polymers were only observed at a concentration of MreB above 0.55 μM (0.02 mg/mL) (Fig. 2B and Table S2).

**Figure 2.**
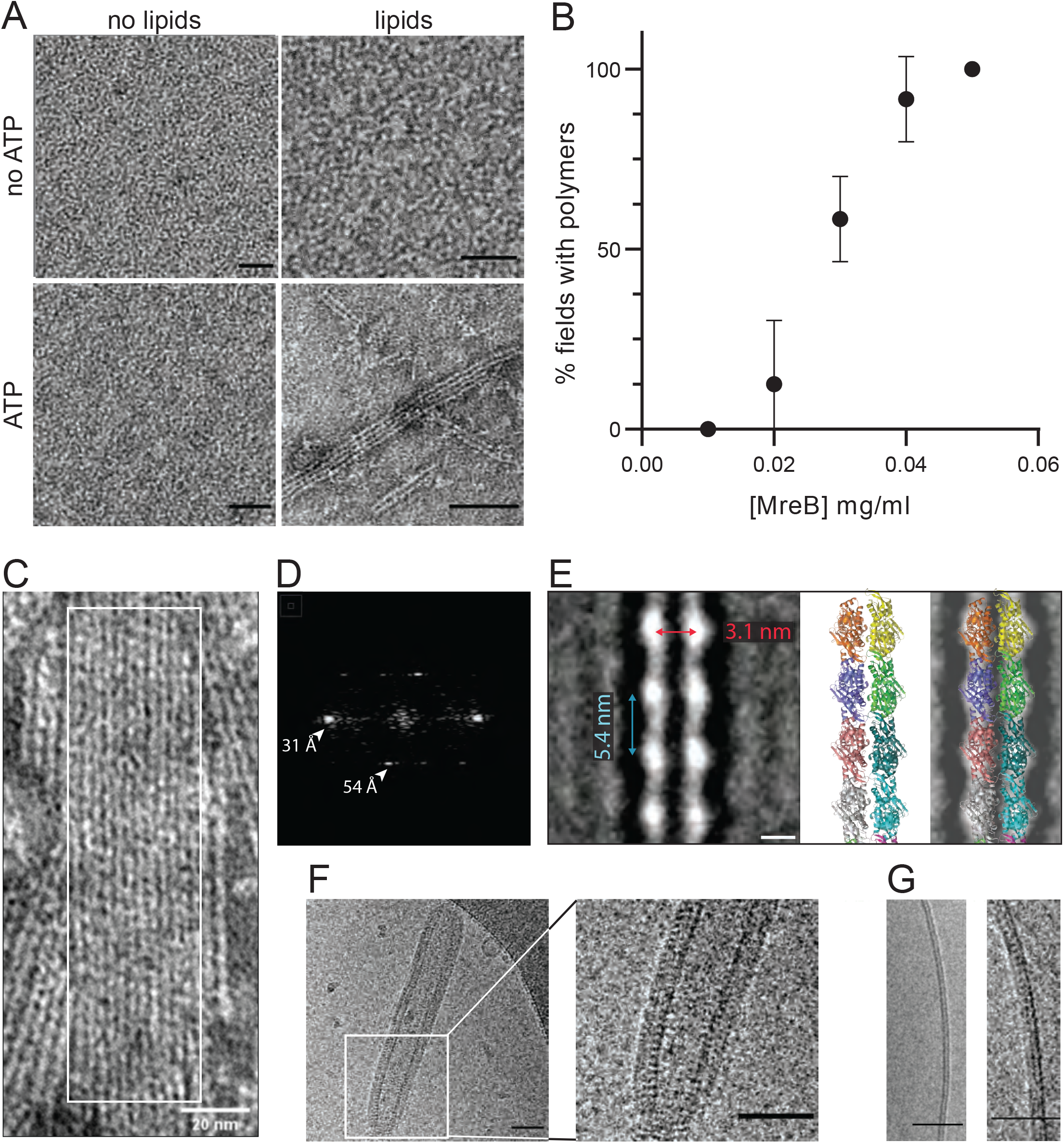
MreB^Gs^ forms double protofilaments in the presence ATP and lipids. (**A**) Polymerization of MreB^Gs^ depends on the presence of lipids and ATP. Negative stained TEM images of purified MreB^Gs^ (0.05 mg/mL) in the presence or absence of 0.5 mg/mL lipid total extract from *E. coli* and of 2 mM ATP. Scale bars, 50 nm. (**B**) Polymer formation as a function of MreB^Gs^ concentration. MreB^Gs^ was set to polymerize in standard conditions at a concentration ranging from 0.01 to 0.05 mg/mL. Values are average of two independent experiments. (**C, D**) MreB^Gs^ polymers assemble into sheets (C). Fourier transform (D) was obtained from the area indicated by a white box in C and revealed a longitudinal subunit repeat of the filaments of 54 Å and a lateral spacing of 31 Å (arrowheads). (**E**) (*Left*) 2D averaging of images of negatively stained dual protofilaments of MreB^Gs^ from 1 554 individual particles. Scale bar, 3 nm. Two copies of the atomic structure of the protofilaments found in the MreB^Gs^ crystals shown to scale (*Middle*, for illustration the two protofilaments are displayed in an antiparallel conformation) and docked into the 2D averaged EM image (*Right*). (**F**) MreB^Gs^ polymers assemble on lipid bilayers and distort liposomes as shown by cryo-electron microscopy (cryo-EM). Cryo-EM micrographs of liposomes (0.37 mg/mL) made from *E. coli* lipid total extracts incubated with purified MreB^Gs^ (0.05 mg/mL) in the presence of ATP (2 mM). Scale bars, 50 nm. (**G**). Cryo-EM micrographs showing the cross-section of the membrane of liposomes in the absence (*Left*) and in the presence (*Right*) of MreB^Gs^. Scale bars, 50 nm.

The simplest assemblies were paired protofilaments, as observed for MreB^Tm^ both in the presence and in the absence of lipids (Salje *et al*., 2011), for MreB^Cc^ assembled on lipid monolayers (van den Ent *et al*., 2014) and for MreB5^Sc^ in solution (Pande *et al*., 2022). Pairs of MreB^Gs^ filaments are generally straight, and individual protofilaments were never observed. Paired protofilaments of different lengths, ranging from below 50 nm up to several micrometers, as well as two-dimensional sheets of straight dual protofilaments could be observed on the same EM grid (Fig. 2A and C, and Fig. S3). In addition, pairs of filaments and sheets always lay flat, indicating that they are oriented relative to the membrane surface. The diffraction patterns of the sheets showed a longitudinal repeat of 54 Å and a lateral spacing of 31 Å (Fig. 2C and D). 2D averaging of negatively stained EM images of 1 554 individual pairs of filaments (Fig. 2E and Fig. S4) also displayed a longitudinal subunit repeat of 54 Å and a lateral subunit repeat of 31 Å, and could well accommodate two scaled protofilaments found in the MreB^Gs^ crystals (Fig. 2E). However, it is not possible to derive the orientation of the protofilaments from the EM density obtained from 2D averaging.

MreB^Gs^ filaments also formed on lipid bilayers as observed by cryo-electron microscopy (cryo-EM). To this end, we prepared liposomes from *E. coli* lipid total extract, and incubated them with MreB^Gs^ and ATP. Lipid vesicles alone were spherical (Fig. S5A), but vesicles decorated with MreB^Gs^ filaments appeared strongly deformed, forming faceted and tubular structures (Fig. 2F and Fig. S5B). These deformed vesicles confirmed that MreB^G^ was bound to the membrane. MreB^Gs^ largely coated the liposomes and displayed a regular pattern along the cross-section of the tubulated vesicles (Fig. 2F and G). This pattern is compatible with longitudinal sections of 2D-sheets of straight filaments aligned in parallel to the longitudinal axis of the cylinder, as previously suggested for the arrangement of MreB^Tm^ in rigid lipid tubes (van den Ent *et al*., 2014).

### ATP or GTP hydrolysis is required for MreB^Gs^ polymerization

In actin, ATP binding or hydrolysis are not required for polymerization (De La Cruz *et al*, 2000; Kasai *et al*, 1965). ATP hydrolysis only occurs subsequent to the polymerization reaction, destabilizing the filaments upon release of the γ-phosphate (Korn, 1982; Korn *et al*, 1987). In contrast, MreB^Tm^ was reported to require either ATP or GTP to polymerize (Esue *et al*., 2006; Nurse & Marians, 2013; van den Ent *et al*., 2001). MreB from *E. coli, C. crescentus, S. citri* and *Leptospira interrogans* also formed polymers in the presence of ATP, but the requirement of ATP for polymerization was not clearly established (Barko *et al*, 2016; Harne *et al*., 2020; Maeda *et al*., 2012; Nurse & Marians, 2013; Salje *et al*., 2011; van den Ent *et al*., 2014). Filaments or sheets of filaments were also observed in the presence of ADP (Gaballah *et al*., 2011; Pande *et al*., 2022; Popp *et al*., 2010b) or AMP-PNP (Pande *et al*., 2022; Salje *et al*., 2011).

Next, we wondered about the specificity of MreB^Gs^ toward nucleotides and their role in the polymerization cycle. MreB^Gs^ formed straight pairs of protofilaments and sheets in the presence of either ATP or GTP, as shown by negative stain EM (Fig. 3A). Noteworthy, the average length of double filaments was increased in the presence of GTP compared to ATP (Fig. S6A), which may reflect differential affinity, dissociation rate or hydrolytic activity of the two nucleotide triphosphates (NTPs). Next, we asked whether the nucleotides diphosphate and monophosphate could also support polymer assembly. As shown in Figure 3A, neither ADP nor GDP or AMP supported filament formation, suggesting that binding and/or hydrolysis of NTPs is required for MreB^Gs^ filament assembly on the lipid monolayer. To discriminate between ATP binding and ATP hydrolysis, we used the non-hydrolysable ATP analogues AMP-PNP and ApCpp (5’-adenylyl methylenediphosphate). No filaments were detected in the presence of either AMP-PNP or ApCpp (Fig. 3A), suggesting that NTP hydrolysis triggers MreB^Gs^ polymerization. However, differential affinity of MreB for these nucleotides could also explain these results. Both actin (Cooke & Murdoch, 1973; Iyengar & Weber, 1964; Kinosian *et al*, 1993) and MreB^Cc^ (van den Ent *et al*., 2014) have the highest affinity for ATP, followed by ADP and then by AMP-PNP. To exclude that the absence of polymerization was due to reduced nucleotide binding, we first increased the concentration of ADP and AMP-PNP from 2 mM to 50 mM. Again, no polymers were detected in the negatively stained samples (Fig. S6B). Next, we performed a competition experiment by mixing ATP (1mM) with increasing amounts of AMP-PNP (1, 10 and 25 mM) in the polymerization reaction. Increasing amounts of AMP-PNP efficiently decreased the presence of MreB^Gs^ filaments on the EM grids (Fig. 3B), indicating that AMP-PNP binds to MreB^Gs^ but does not support efficient polymerization. Taken together, these results suggest that ATP hydrolysis is required for assembly of MreB^Gs^ into filaments on a membrane surface.

**Figure 3.**
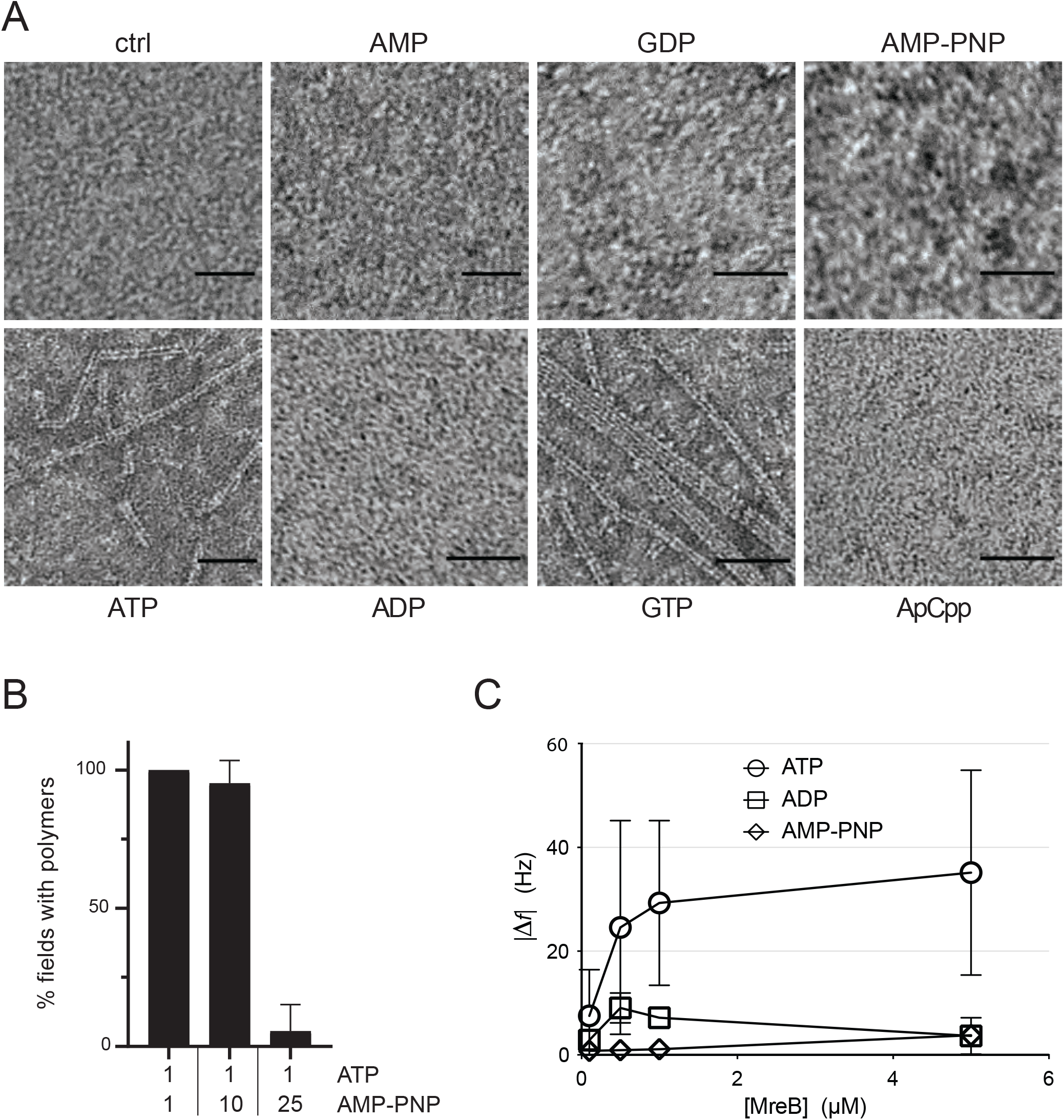
Polymerization of *Geobacillus* MreB depends on the presence of hydrolysable nucleotides. (**A**) ATP and GTP promote assembly of MreB^Gs^ polymers. Negative stained EM images of purified MreB^Gs^ (0,05 mg/mL) incubated in the presence of either ATP, ADP, AMP, GTP, GDP, the non-hydrolysable AMP-PNP or ApCpp (2 mM), or in the absence of nucleotide (ctrl) on a lipid monolayer, for 2 h at room temperature. Scale bars, 50 nm. (**B**) Formation of MreB^Gs^ double filaments on a lipid monolayer depends on ATP hydrolysis. AMP-PNP competes with ATP for binding to MreB^Gs^, preventing polymerization. MreB^Gs^ was set to polymerize in standard conditions except that 2 mM ATP was replaced by a mix of ATP and AMP-PNP at the indicated concentrations (in mM). Values are average of three independent experiments. (**C**) Adsorption of MreB^Gs^ to a supported lipid bilayer depends on ATP hydrolysis. Frequency changes (|Δ*f*|) in QCM-D experiments measured with varying amount (0.1 - 5 μM) of MreB^Gs^ on a SLBs made of DOPC:DOPG 80:20 and in the presence of 2 mM of either ATP, ADP or AMP-PNP.

### Nucleotide hydrolysis is required for binding of MreB^Gs^ to the membrane, as monolayered MreB films

We next wondered whether NTP hydrolysis triggers the binding of MreB^Gs^ monomers to the membrane prior to polymerization or whether it promotes the polymerization of membrane-bound monomers. To address this question, we turned to quartz crystal microbalance with dissipation monitoring (QCM-D) to measure the binding affinity of MreB^Gs^ to supported lipid bilayers (SLBs) of various lipid mixtures. QCM-D is a surface-sensitive technique that can be used to measure biomolecular interactions at aqueous interfaces in real time (Reviakine *et al*, 2011). Changes in frequency (Δ*f*) and dissipation (Δ*D*) are recorded. The frequency is directly proportional to any mass added or removed (Sauerbrey, 1959), while dissipation changes are indicative of the viscoelastic properties of the attached layer. QCM-D was previously applied to study, for example, the binding affinity of the division proteins MinD and MinE of *E. coli* to SLBs (Renner & Weibel, 2012). *E. coli* and *B. subtilis* membranes are mainly composed of phospholipids, with the anionic phosphatidylglycerol (PG) and the zwitterionic phosphatidylethanolamine (PE) being the dominant species (Bernat *et al*, 2016; Bishop *et al*, 1967; den Kamp *et al*, 1969; Laydevant *et al*, 2022; Nickels *et al*, 2017; Seydlova & Svobodova, 2008; Sohlenkamp & Geiger, 2016). Although lipid proportions vary widely depending on the strains and growth conditions, PE is largely dominant in *E. coli* while PG is more dominant in *B. subtilis*, indicating that phospholipids are more negatively charged in G+ membranes. To mimic *Bacillus* membranes in our QCM-D assay, we used mixtures of the zwitterionic dioleoylphosphatidylcholine (DOPC) doped with the anionic dioleoylphosphatidylglycerol (DOPG) in different proportions (100% DOPC, 90:10 DOPC:DOPG, 80:20 DOPC:DOPG) to generate SLBs. DOPC was selected to replace PE because of its widespread role as a scaffold lipid in SLBs formation. We had to adopt a mixture that enabled us to form SLBs on planar substrates, as the inverted conical shape of PE makes the formation of planar SLBs difficult (PE has a tendency to form non-bilayer structures because of its small headgroup). A typical SLBs signature experiment is shown in Fig. S7A-B. Briefly, SLBs are formed after the adsorption of liposomes (Δ*f* decrease, Δ*D* increase) onto activated silica surfaces. Once a critical surface concentration of liposomes is reached and the interactions between liposomes and the surface are suitable, the liposomes spontaneously rupture and coalesce into flat SLBs (Keller *et al*, 2000). After the formation of stable and flat SLBs (i.e. a stable baseline for frequency and dissipation) (Fig. S7A), we started to add MreB^Gs^ to the SLBs (Fig. S7B, closed arrows). We recorded frequency and dissipation changes for the added MreB^Gs^ protein (in varying concentrations ±2mM ATP) on all SLBs. Binding was strongly dependent on ATP (Fig. 3C and S7C-D) and was substantially affected by the lipid composition of SLBs (Fig. S7C). Increasing the levels of DOPG led to a higher amount of MreB^Gs^ binding, with DOPC:PG 80:20 giving the highest observed adsorption, suggesting that the presence of negatively charged lipids favors MreB^Gs^ binding. Binding was detected almost instantaneously after adding MreB^Gs^ (Fig. S7B, closed arrows) for all concentrations of MreB tested, either above or below the concentration in which polymers were observed by EM (0.55 μM, Fig. 2B and Table S2). The protein binding kinetics reached an equilibrium after approximately 5-10 min with a somewhat slower continued binding of additional MreB^Gs^ monomers (Fig. S7B). These observations suggested that in the presence of ATP both monomers and polymers of MreB^Gs^ can interact with the membrane. However, upon rinsing with the same buffer (Fig. S7B, open arrows), MreB^Gs^ at low (monomeric) concentrations was completely removed from the membrane while polymeric MreB^Gs^ remained more stably absorbed. When replacing ATP with ADP or AMP-PNP, we were not able to detect any significant binding, indicating a virtually complete loss of interaction (Fig. 3C). We further increased the concentration of ADP or AMP-PNP to exclude the possibility that the binding was simply affected by a decreased affinity of MreB^Gs^ for these nucleotides. Higher concentrations of ADP and AMP-PNP did not restore the binding of MreB^Gs^ to the SLBs (Fig. S7D). We concluded that nucleotide hydrolysis provides the energy required for MreB^Gs^ membrane binding and that filaments bind more stably than MreB^Gs^ monomers.

Finally, we used the Sauerbrey model (Sauerbrey, 1959) to calculate the average coverage and thickness of the layer of MreB^Gs^ attached to the SLB. The thickness of the MreB films ranged from 0.1 nm to approximately 4 nm on the SLBs with a ratio of DOPC:DOPG 80:20, which corresponds to ~ 2.5% to 100% coverage assuming a monolayer filament thickness (Fig. S7E and Material and Methods). These data suggest that MreB^Gs^ mainly form monolayers on the SLBs, with limited out-of-plane interactions (i.e. limited tendency to stack into multilayers), consistent with our EM observations of pairs of filaments and sheets lying flat on the lipid monolayer (Fig. 2, Fig. 3 and Fig. S3), with the pattern displayed by the filaments on cross-sections of vesicles (Fig. 2G), and thus with the interaction of the membrane with a specific surface of the MreB^Gs^ filaments. Taken together, these observations suggest an oriented arrangement of MreB^Gs^ filaments on the membrane, with lateral interactions between filaments in the plane perpendicular to their membrane-binding surface.

### The amino-terminus and the α2β7 hydrophobic loop of MreB^Gs^ are required for membrane binding and polymerization

In MreB^Tm^, membrane-binding is mediated by a small loop containing two hydrophobic residues (L93 and F94), whereas in MreB^Ec^ and MreB^Cc^ it is mediated by an amino terminal extension (~9 residues) predicted as an amphipathic helix, which is disordered in all crystal structures of MreB^Cc^ (Salje *et al*.,2011; van den Ent *et al*., 2014), (Fig. S1, green highlights). Albeit essential to MreB function in *E. coli* (Salje *et al*., 2011), this N-terminal extension is not required for polymerization *in vitro* (Salje *et al*., 2011; van den Ent *et al*., 2014). MreB^Bs^ was not predicted to carry an N-terminal amphipathic helix (Salje *et al*., 2011). A systematic search in a large panel of MreB proteins spanning over the entire bacterial kingdom revelaed that N-terminal amphipathic helices are a conserved feature of the Proteobacteria phylum and most G- bacteria, but are absent from *Firmicutes* and *Bacteroidetes* species (Fig. S8). Most *Firmicutes*, including *Bacilli* (MreB^Gs^ and MreB^Bs^) and *Clostridia* but to the notable exception of the wall-less *Mollicutes* (or, put it in other words, G+ bacteria to the exception of *Actinobacteria*) possess a shorter N-terminal sequence containing 4-7 hydrophobic amino-acids (Fig. S1 and Fig. S8). We noticed that in the crystal structure of the apo form of MreB^Gs^ this short hydrophobic N-terminal sequence is in close proximity to loop α2-β7 (Fig. 3A), which in MreB^Tm^ carries the hydrophobic residues L93 and F94 involved in membrane binding (Salje *et al*., 2011). The α2-β7 loops of MreB^Bs^ and MreB^Gs^ contain additional hydrophobic residues (Fig. S1), suggesting that they may also play a role in membrane interaction. We constructed and purified mutants deleted for either four hydrophobic residues of the α2-β7 loop (aa 95-98, GLFA), the N-terminal sequence 2-7 (FGIGTK), or both (Table S3). Folding of the protein was not affected by the deletions as shown by circular dichroism (CD) (Fig. S9). The three mutants and the wild-type MreB^Gs^ protein were set to polymerize in our standard conditions and the formation of filaments was assessed by negative stain EM. The three mutants displayed a dramatic reduction of their polymerization capabilities with a gradation of defects, the deletion of the N-terminal sequence having the lowest impact and the double deletion the highest (Fig. 4A and Fig. S10).

**Figure 4.**
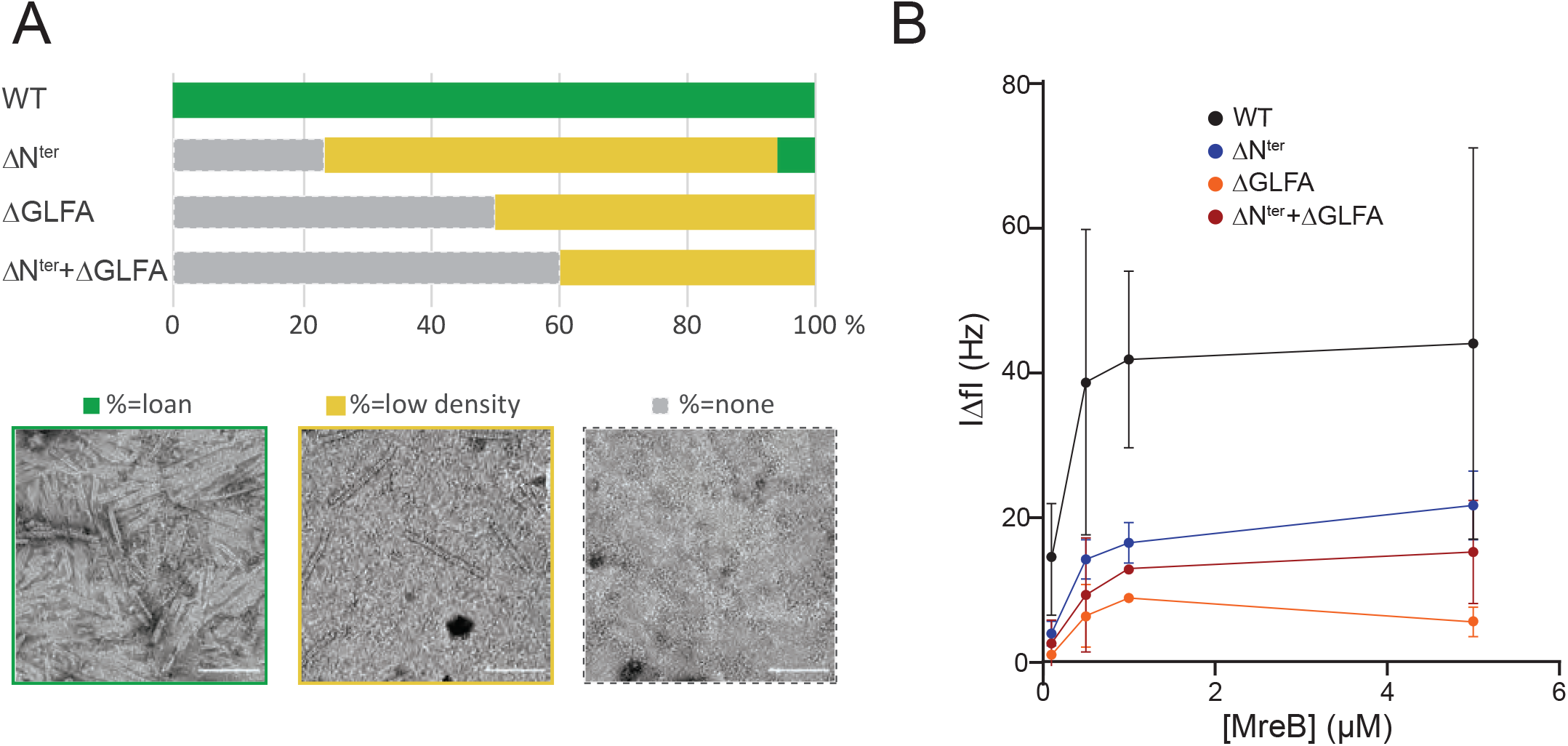
The N-terminus and the α2β7 hydrophobic loop of MreB^Gs^ promote membrane binding and polymerization. (**A**) Both the hydrophobic α2-β7 loop and the N-terminus sequence of MreB^Gs^ are required for efficient polymerization on a lipid monolayer. Frequency and density of polymer formation observed on negatively stained TEM images for the wild type (WT) and the mutants of the α2-β7 (ΔGLFA), the N-terminus (ΔN^ter^) or both domains (ΔN^ter^+ΔGLFA) of MreB^Gs^. Images were categorized based on absence or the presence of low or high density of polymers. Values are the sum of 2 independent experiments. (**B**) The α2-β7 loop and the N-terminus domain of MreB^Gs^ enhance its adsorption to supported lipid bilayers. Frequency change (|Δ*f*|) measured for the binding of various concentrations (0.1 - 5 μM) of purified wild-type (WT) and mutant forms of MreB^Gs^ to SLBs. Incubations were performed in polymerization buffer containing 2 mM ATP. SLBs contained an 80:20 molecular ratio of DOPC:DOPG.

We next tested whether the polymerization defect observed with the mutants was due to a lack of interaction with the lipids, as expected. In QCM-D experiments, membrane adsorption in the presence of ATP was strongly reduced in the three mutants relative to the wild-type protein (Fig. 4B), mirroring the polymerization assays (Fig. 4A). As expected, in the presence of ADP, binding was not observed for any of the mutants, as observed with the wild-type protein (Fig. S7F). Taken together, these results suggest that the spatially close hydrophobic N-terminus and α2-β7 loop are the membrane anchors of MreB^Gs^. Deletion of these hydrophobic motifs prevents MreB^Gs^ ATP-dependent binding to lipids, which in turn prevents filament formation.

### γ-phosphate dissociation after ATP/GTP hydrolysis by MreB^Gs^ is related to filament turnover

Our results indicate that MreB^Gs^ has a limited intrinsic affinity for lipids, with nucleotide hydrolysis switching the protein from a soluble to lipid-affine form, potentially through a structural change. In order to test the impact of nucleotide binding, we co-crystallized MreB^Gs^ with ATP-Mg and solved the crystal structures of the complex at 2.3 Å resolution (PDB ID 8AZG). The crystals diffracted in space group P2_1_2_1_2 (Table S1) and the analysis of the packing did not reveals the formation of protofilaments. The structure of the ATP-bound form of MreB^Gs^ is highly similar to the apo form of the protein, with a rmsd of 1.41 Å over 313 aligned Cα atoms (Fig. S11A). However, ATP binding induces a small closure of the nucleotide-binding pocket, and loop β6-α2, which was disordered in the apo structure, is now fully visible in the electron density map. The hydrophobic loop α2-β7 and the N-terminus also display an alternative conformation. Interestingly, despite highly conserved nucleotide-binding residues, the γ-phosphate of the bound ATP occupies the position of the Mg ion observed in the crystal structure of MreB^Cc^ bound to the non-hydrolysable ATP analog AMP-PNP (PDB ID 4CZJ) (Fig. S11B and C). Despite multiple co-crystallization trials, crystal packing never revealed straight protofilaments like in the apo structure, only monomers were present in the ATP-bound state.

MreB of several G- bacteria was previously shown to slowly hydrolyze ATP in solution (Bean & Amann, 2008; Esue *et al*., 2005; Esue *et al*., 2006; Gaballah *et al*., 2011; Mayer & Amann, 2009; Nurse & Marians, 2013; Pande *et al*., 2022; Popp *et al*., 2010b). Our QCM-D results suggested that ATP hydrolysis by MreB^Gs^ is a prerequisite for membrane binding and polymerization, and that it may thus occur in solution too. We monitored ATPase activity by measuring the release of inorganic phosphate (P_i_) in the presence of ATP for a wide range of MreB concentrations, in the presence and in the absence of lipids. In the absence of lipids, the equilibrium rate of P_i_ dissociation was 0.032 ± 0.002 P_i_/min/MreB molecule at 37°C, and 0.081 ± 0.004 P_i_/min/MreB at 53°C, a temperature closer to the optimal growth temperature of *G. stearothermophilus* (Fig. 5A and Fig. S12A). In the presence of lipids, the rate of P_i_ release increased ~2-fold, to 0.065 ± 0.005 P_i_/min/MreB at 37°C and 0.158 ± 0.003 P_i_/min/MreB at 53°C (Fig. 5A and Fig. S12A). These rates of P_i_ release upon ATP hydrolysis (~ 1 P_i_/MreB in 6 min at 53°C) are comparable to those observed for MreB^Tm^ and MreB^Ec^ *in vitro* (Esue *et al*., 2005; Esue *et al*., 2006; Nurse & Marians, 2013), and also remarkably similar to those of the (very slow) dissociation of γ-phosphate after ATP hydrolysis within actin filaments, which has a half-time of ~6 min (dissociation rate constant ~ 0.003 sec^-1^) (Carlier & Pantaloni, 1986). Interestingly, the release of P_i_ was constant for hours, over the length of our ATPase experiments (Fig. 5B). However, similar density and lengths of negatively stained MreB^Gs^ polymers were observed over the EM grids for all incubation (polymerization) times tested, ranging from a few minutes to several hours (Table S2). These observations suggest that MreB polymerization in the presence of lipids is a dynamic process, with steady state polymerization/depolymerization rates.

**Figure 5.**
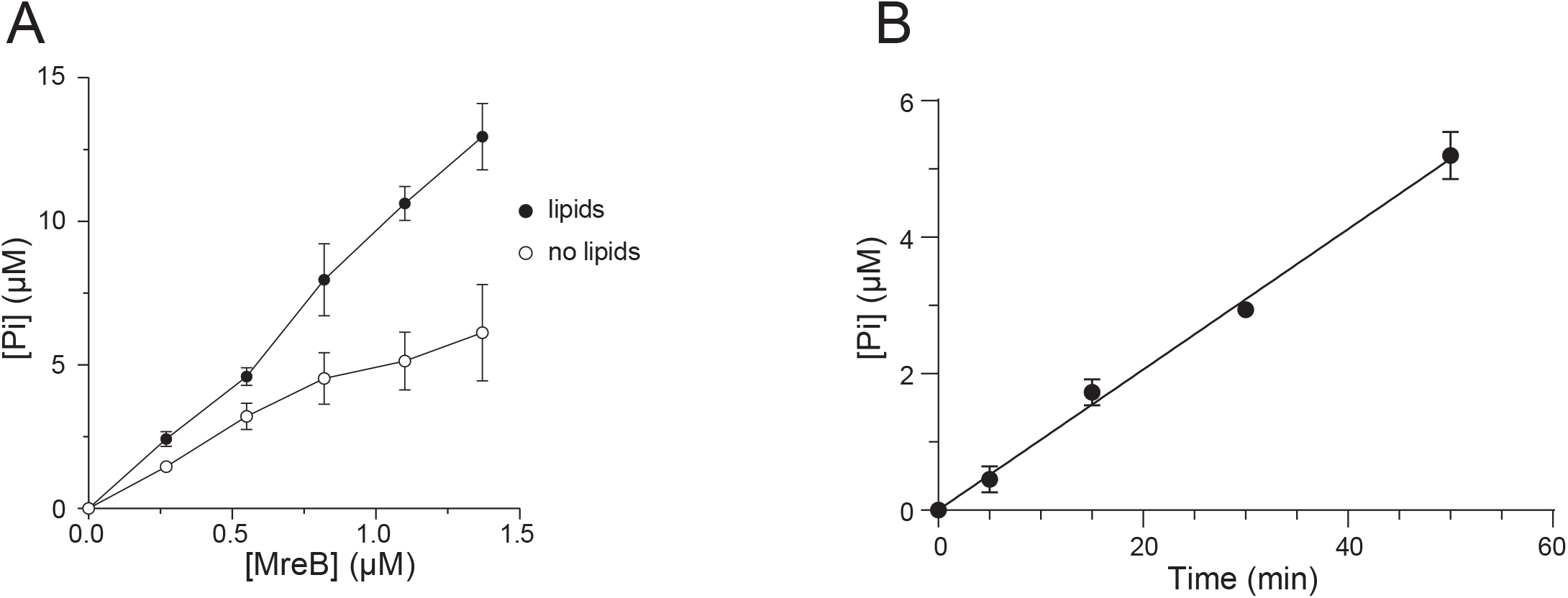
ATPase activity of MreB^Gs^. (**A**) The ATPase activity of MreB^Gs^ is stimulated in the presence of lipids. ATPase activity, measured by monitoring inorganic phosphate (Pi) release, of MreB^Gs^ at different concentrations (0.27 – 1.37 μM) in the presence of 0.5 mM ATP and in the presence or absence of 0.05 mg/mL liposomes, after 1 h incubation at 53°C. Values are averages of at least 2 independent experiments. Error bars are standard deviations. (**B**) Kinetics of ATP hydrolysis detected via P_i_ release in the presence of 1.37 μM MreB^Gs^, 0.5 mM ATP and 0.05 mg/mL liposomes at 53°C. The line is a simple linear regression fit (goodness of fit R^2^ = 0.9910). Values are averages of 2 independent experiments. Error bars are standard deviations.

Finally, we have shown that, in solution, MreB^Gs^ polymerizes in the presence of lipids and either ATP or GTP (Fig. 3A and Fig. S6A). MreB^Tm^ was reported to polymerize in solution in the presence of ATP or GTP as well (Bean & Amann, 2008; Esue *et al*., 2006; Nurse & Marians, 2013; Popp *et al*, 2010a; van den Ent *et al*., 2001), and to release P_i_ at similar rates upon GTP and ATP hydrolysis (Esue *et al*., 2006). We found that MreB^Gs^ also releases P_i_ after hydrolysis of GTP as efficiently as after hydrolysis of ATP, both in the presence and in the absence of lipids (Fig. S12B).

Taken together, these results indicate that the presence of lipids is not required for the ATPase/GTPase activity of MreB^Gs^. However, the presence of lipids stimulates P_i_ release, advocating for some conformational changes upon MreB^Gs^ binding to the lipids and/or upon polymerization on the lipid surface. Furthermore, P_i_ release is slow but constant over extended periods of time while filament length and density remain unchanged, suggesting a dynamic filament assembly/disassembly process.

## Discussion

Here we show that bacterial actin MreB from the G+ bacterium *G. stearothermophilus* polymerizes into pairs of protofilaments on lipid membranes. In contrast to G- MreBs, which were shown to also polymerize in bulk solution, polymerization of MreB^Gs^ was only observed in the presence of lipids. The requirement of the membrane for polymerization is consistent with the observation that MreB polymeric assemblies *in vivo* are membrane-associated only (i.e. localize at the cell periphery but not in the cytoplasm), in line with their role as scaffold of the CW elongation machinery. Membrane binding of MreB^Gs^ is direct and mediated by the hydrophobic α2-β7 loop protruding from the protein in domain IA, in line with the prediction of Salje and colleagues that binding to membranes via such hydrophobic loop and/or an amphipathic helix may be conserved for all MreBs (Salje *et al*., 2011). However, we found that binding of MreB^Gs^ to the membrane is also mediated by the hydrophobic N-terminus which, together with the spatially closed α2-β7 loop, would constitute a membrane anchor. The absence of an amphipathic helix and the presence instead of a hydrophobic N-terminus in many MreB sequences (Fig. S8) suggests that most MreB use one or the other amino-terminal structure to bind to membranes.

Another difference relative to G- MreBs concerns the requirement of NTP hydrolysis for membrane binding and polymerization of MreB^Gs^. It was reported that membrane binding by MreB^Tm^ is not dependent on nucleotide binding or hydrolysis (Salje *et al*., 2011). Furthermore, G- MreBs polymerized in the absence of added lipids (Barko *et al*., 2016; Esue *et al*., 2005; Esue *et al*., 2006; Harne *et al*., 2020; Maeda *et al*., 2012; Nurse & Marians, 2013; Popp *et al*., 2010b; Salje *et al*., 2011; van den Ent *et al*.,2001), indicating that membrane binding is not a prerequisite for their polymerization. As demonstrated here, membrane binding by *Geobacillus* MreB requires not only binding but also hydrolysis of the nucleotide, either ATP or GTP. The role of nucleotides on the polymerization of G- MreBs is somewhat confusing in the literature as it varies significantly between reports, even for MreBs from the same species. ATP was found to be essential for G- MreBs polymerization in some reports (Barko *et al*., 2016; Esue *et al*., 2005; Nurse & Marians, 2013; van den Ent *et al*., 2001) while other reports indicate that polymerization also occurs in the presence of ADP (Bean & Amann, 2008; Gaballah *et al*., 2011; Mayer & Amann, 2009; Pande *et al*., 2022; Popp *et al*., 2010b) or AMP-PNP (Pande *et al*., 2022; Salje *et al*., 2011). Filaments were observed for MreB^Tm^ and MreB5^Sc^ in the presence of AMP-PNP, but polymerization in the presence of ADP was in most cases concluded from light scattering experiments alone, so the possibility that aggregation rather than ordered polymerization occurred in the process cannot be excluded. Differences in the purity of the nucleotide stocks used in these studies could also explain some of the discrepancies. On the basis of our data and the existing literature, we propose that the requirement for ATP (or GTP) hydrolysis for polymerization may be conserved for most MreBs.

Taken together, our data suggest a model (Fig. 6) in which nucleotide hydrolysis by MreB^Gs^ in solution may induce a conformational change that allows the membrane-binding motifs of MreB^Gs^ monomers to interact with the membrane, possibly in an ADP-Pi-MreB state as suggested by the very slow rate of P_i_ release. Comparison of the crystal structures of apo MreB^Gs^ and its ATP-bound form shows that only minor conformational changes occur upon nucleotide-binding, in agreement with what was observed when comparing crystal structures of MreB^Cc^ and MreB5^Sc^ in different nucleotide-bound states (Harne *et al*., 2020; Pande *et al*., 2022; van den Ent *et al*., 2014). This invariability of folding regardless of the bound ligands has also been observed in crystal structures of actin and other members of the actin superfamily (Schuler, 2001). ATP hydrolysis and membrane binding might require small but dynamic structural changes that cannot be observed in crystal structures locked in a conformation imposed by the packing. The absence of protofilaments in the crystal packing of the ATP-MreB^Gs^ complex indicates that the surface of ATP-bound MreB monomers was not prone to interaction despite the very high concentration of protein and the crystal packing forces (which explain filament formation in the crystals of the apo form). It is tempting to speculate that ATP-bound MreB is soluble and that polymerization is linked to structural changes upon ATP hydrolysis, consistent with our finding that NTP hydrolysis is required for MreB^Gs^ polymerization. Membrane interaction upon nucleotide hydrolysis would promote polymerization, possibly through a second conformational change (Fig. 6). This second conformational change may favor P_i_ release since the release rate increased 2-fold in the presence of lipids. The rate of P_i_ release from MreB^Gs^ filaments remained nevertheless low, consistent with previous reports on MreB^Tm^, MreB^Ec^ and MreB5^Sc^ (Bean & Amann, 2008; Esue *et al*., 2005; Esue *et al*., 2006; Nurse & Marians, 2013) and was strikingly similar to that from filamentous actin, where the P_i_ release half-time (6 min) is much slower than the ATP hydrolysis half-time (~2 sec) (Pollard, 2016). Thus, for both MreB and actin, despite hydrolyzing ATP before and after polymerization, respectively, the ADP-Pi-MreB intermediate would be the long-lived intermediate state within the filaments. In actin, the release of γ-phosphates after ATP hydrolysis within the filaments induces a conformational change that destabilizes the filament and promotes depolymerization. Importantly, the release of the γ-phosphate by MreB^Gs^ in polymerization conditions continued well after steady-state levels of polymerization were achieved (Fig. 5B). Two scenarios could explain this: (i) a constant but extremely slow release of P_i_ from stable filaments or (ii) a turnover of the filaments. We hypothesize that MreB filaments turnover and that, as in actin, the release of P_i_ is involved in this process.

**Figure 6.**
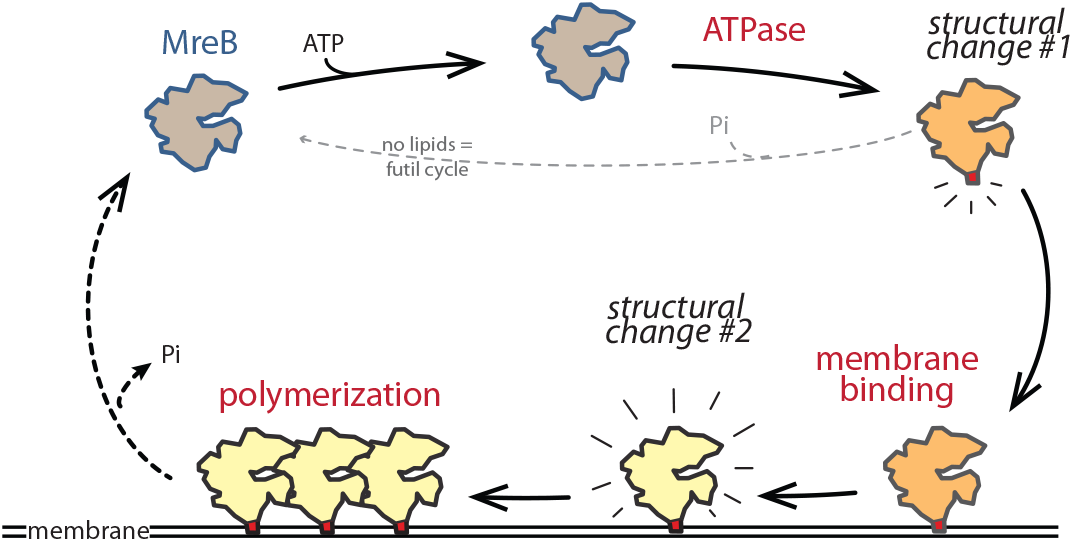
Model for ATPase-dependent membrane binding and polymerization of MreB^Gs^. MreB polymerization follows a hierarchy of events. ATP hydrolysis by monomeric MreB^Gs^ (grey) stimulates MreB^Gs^ adsorption to lipids, possibly by promoting a conformational change (orange) that renders the α2-β7 loop and N-terminal sequence (red motif) prone for interaction with the membrane MreB^Gs^ monomers competent for lipid interaction, possibly in the ADP-P_i_ form, form then membrane-associated polymers. The absence of polymers in the presence of ATP and absence of lipids supports a model in which a second conformational change (light yellow) may occur upon binding of MreB monomers to the membrane, which triggers polymerization.

Our EM and cryo-EM data show that MreB^Gs^ filaments are straight and therefore most likely rather rigid. In agreement with this hypothesis, lipid vesicles coated with MreB^Gs^ filaments were strongly deformed and faceted (Fig. 2E and Fig. S5B). However, MreB^Gs^ filaments outside liposomes did not bend the liposomes into negative curves as previously reported for MreB^Tm^ and MreB^Cc^ (Salje *et al*., 2011; van den Ent *et al*., 2014). A recent model postulates that MreB polymers are intrinsically curved and have affinity for negatively curved membranes while avoiding to be positively bent (Hussain *et al*., 2018; Wong *et al*, 2019). The pattern of MreB^Gs^ filaments in longitudinal sections of coated tubulated liposomes (Fig. 2G) is compatible with straight filaments aligning with the longitudinal axis of the rod to avoid positive curvature. However, the intrinsic affinity of MreB filaments for negative concave membrane curvature remains however to be conclusively demonstrated. The kinetics of polymerization of MreBs, as well as the presumed apolarity of growth of antiparallel doublets, are also questions for future studies.

## Materials and Methods

### General procedures and growth conditions

DNA manipulations were carried out by standard methods. *G. stearothermophilus* was grown at 59°C and *E. coli* at 37°C in rich lysogeny broth (LB). Kanamycin was used at 25 μg/mL. All strains used in this study are listed in Table S4. All lipids, *E. coli* Lipid Total Extract (TE), 1,2-Dioleoyl-sn-glycero-3-phosphocholine (DOPC), and 1,2-dioleoyl-sn-glycero-3-phospho-(1’-rac-glycerol) (DOPG), were purchased from Avanti Polar Lipids Inc. (Alabaster, AL).

### Cloning, expression and purification of MreB variants from *G. stearothermophilus*

Two *mreB* paralogs were identified in the genome of *G. stearothermophilus* ATCC 7953, corresponding to *mreB* and *mbl* of *B. subtilis* based on their synteny. The *mreB* ortholog displays a strong 92.6 % similarity (85.6 % overall identity) with *mreB* of *B. subtilis* (Fig. S1). *mreB* from *G. stearothermophilus* ATCC 7953 was amplified by PCR using primers cc430 and cc431 (Table S5) and *G. stearothermophilus* growing cells as template. A second DNA fragment was generated by PCR on a derivative of plasmid pET28a (devoid of the first three codons following the *NcoI* restriction site), using primers cc433/cc432 (Table S5). The two resulting fragments were assembled by isothermal assembly and transformed into *E. coli* (Gibson *et al*, 2009). The resulting plasmid, pCC110, which carries a wild-type version of *mreB^Gs^* in translational fusion with a 5’ extension encoding a 6-histidine tag, was used as a template to generate pCC116, pCC117 and pCC115, carrying the *mreB^Gs^* gene deleted for codons 2-7 (FGIGTK), 102-105 (GLFA), or both, respectively. For this, pCC110 was PCR-amplified using primers cc582/cc583 (to generate pCC116) or cc584/cc585 (to generate pCC117) (Table S5) and the PCR products were treated with *Dpn*I prior to transformation into *E. coli*. To generate pCC115, isothermal assembly was performed with two PCR products generated using primers cc582/cc585 and cc583/cc584 and pCC110 as template, and the product was transformed into *E. coli*. Following extraction and sequencing, the four resulting pCC plasmids were transformed into the T7 *express E. coli* expression host (Table. S4).

The his-tagged proteins were produced in T7 express *E. coli* cells grown in LB broth supplemented with kanamycin. Expression of recombinant MreB was induced by the addition of IPTG at the final concentration of 1 mM, when cultures reached an optical density at 600 nm of 0.6. Expression was performed over-night at 15°C, with maximum aeration. Bacteria were harvested by centrifugation (5 000 *g* for 7 min at 4°C).

Pellets were resuspended in buffer A (20 mM Tris pH 7, 500 mM KCl) supplemented with EDTA-free complete protease inhibitor (Roche) and 250 μg/mL of lysozyme. Cells were disrupted by sonication on a Vibra-Cell VC505 processor (Sonics & Materials, Inc., Newton, CT, USA) for 10 min with 10 seconds on/off cycles at 50% power, and the supernatant was collected after clarification by centrifugation (40 000 *g* for 20 min at 4 °C). The 6-histidine-tagged MreB variants followed a two-step purification procedure. The proteins were first purified by affinity chromatography on a Ni-nitrilotriacetic acid (Ni-NTA) agarose resin (Thermo fisher scientific). The column was washed with buffer A supplemented with 20 mM imidazole, and proteins were eluted with a step gradient of imidazole (100 mM to 400 mM) in buffer A. The collected fractions were analyzed by electrophoresis, using a 12% polyacrylamide precast gel (Mini-PROTEAN TGX stain free, Bio-Rad). Fractions containing the purest form of the proteins were loaded on a size exclusion chromatography HiLoad^®^ 16/60 Superdex^®^ 200 pg column (GE Healthcare Life Sciences / Cytiva), pre-equilibrated with buffer B (buffer A supplemented with 1 mM DTT and 2 mM EDTA) connected to an AKTA FPLC system (GE Healthcare Life Sciences). Fractions corresponding to the elution peaks were analyzed by electrophoresis to assess the presence of MreB, pooled and concentrated with an ultrafiltration spin column (Vivaspin, 10 000 MWCO), up to a maximum of 0.5 mg/mL (14 μM), as determined from the absorption at 280 nm measured using a Nanodrop spectrophotometer (Thermo fisher scientific). The recombinant proteins were aliquoted and immediately frozen and stored at −20°C.

### Preparation of lipid monolayers and negative stain electron microscopy

MreB was set to polymerize for 2-3 hours (unless stated otherwise, Table S2) at room temperature in the reaction buffer (4-20 mM Tris pH7, 100-500 mM KCl, 1-5 mM Mg^2+^) with or without a lipid extract and with or without nucleotide (Fig. 2A, Fig. S2A and Table S2).

Polymerization of MreB on lipids was induced by creating a lipid monolayer on droplets containing MreB (typically 0.05 mg/mL) in the reaction buffer. Lipids from *E. coli* TE were dissolved to 2 mg/mL in chloroform in a glass vial and stored at −20°C. Lipid preparations were diluted in chloroform to a final concentration of 0.5 mg/mL on the day of the experiment. Approximately 200 nL of lipid preparation were dropped on top of the droplets containing MreB in the reaction buffer, previously placed on a solid support in a humid chamber, and incubated at room temperature. The standard reaction buffer supporting polymerization contained 20 mM Tris pH7, 500 mM KCl, 2 mM ATP and 5 mM Mg^2^.

For TEM observations, a carbon-coated electron microscopy grid (CF300-Cu, Electron Microscopy Sciences), carbon side down, was placed on top of lipid-coated reaction droplets and gently lifted after 2 minutes incubation. Grids were stained with either a solution of 2% uranyl formate or 1% uranyl acetate and air-dried prior to TEM observation. TEM images were acquired on a charge-coupled device camera (AMT) on a Hitachi HT 7700 electron microscope operated at 80kV (Milexia – France) or a Tecnai G2 LaB6 (Thermofischer FEI) microscope operated at 200 kV or a Tecnai Spirit (Thermofischer FEI) microscope operated at 80 kV.

Fourier Transformation of MreB sheets was done using ImageJ to obtain diffraction patterns. For 2D processing, a set of images was collected at a magnification of 50 000× with a pixel size of 2.13 Å per pixel and a defocus varying from −2 to −1 μm, using a Tecnai G2 LaB6 (Thermofischer FEI) microscope operated at 200 kV and a F-416 TVIPS 4K×4K camera. To obtain 2D class averages, particles were classified and aligned, using SPIDER (Frank *et al*, 1996). 1 554 Particles were windowed out into 99 × 99 pixels images by using the Boxer interface of EMAN (Ludtke *et al*, 1999) and appended into a single SPIDER file, then normalized against the background. One round of reference-free alignment and classification was performed before references were selected from the first-class averages. Several rounds of multireference alignment and classification were then performed, and new references were selected from the class averages until no further improvement was obtained.

### Quantification of MreB filaments on EM images

We set up a protocol to compare, based on EM images and in a quantitative way, the propensity of MreB to form polymers between different conditions. To circumvent the issue of the highly heterogeneous distribution of polymers on the EM grids that could bias the analysis, we acquired for each experimental replica, images on 12 random locations covering the entire grid. To determine the impact of the concentration of MreB or the nucleotides used on the formation of polymers, images were sorted based on the sole presence or absence of polymers (regardless of their density), and we plotted the % of fields containing MreB filaments. To accurately compare the effect of the ΔN^ter^ and ΔGLFA deletions on MreB ability to polymerize, we refined the classification by sorting the images based on the density of polymers. For this, anonymized images of all strains from two replica were pooled and subsequently distributed based on the density of polymers (none, low density, or loan). 16% of the images were discarded due to low quality. The remaining images were distributed based on the 3 groups and expressed as percentage of fields.

### Preparation of liposomes and cryo-electron microscopy

*E. coli* TE was dissolved in chloroform, aliquoted, dried under a stream of argon, and desiccated for 1 hour under vacuum. The liposome solution was made by resuspending desiccated TE in polymerization buffer (20 mM Tris-HCl pH 7, 500 mM KCl, 2 mM ATP, 5 mM MgCl_2_) on the day of the experiment, to a final lipid concentration of 1 mg/mL. 0.05 mg/mL of purified MreB was mixed with the liposome solution and incubated 2 h at room temperature. 4 μL of sample were applied to a glow-discharged holey lacey carbon-coated cryo-electron microscopy grids (Ted Pella, USA). Most of the solution was blotted away from the grid to leave a thin (<100 nm) film of aqueous solution. The blotting was carried out on the opposite side from the liquid drop and plunge-frozen in liquid ethane at −181 °C using an automated freeze plunging apparatus (EMGP, Leica, Germany). The samples were kept in liquid nitrogen and imaged using a Tecnai G2 (FEI, Eindhoven, Netherlands) microscope operated at 200 kV and equipped with a 4k × 4k CMOS camera (F416, TVIPS). The imaging was performed at a magnification of 50 000× with a pixel size of 2.13 Å using a total dose of 10 electrons per Å^2^.

### Circular dichroism

The secondary structure of recombinant WT and mutant forms of MreB were analyzed by circular dichroism (CD). Far-UV spectra were recorded on a J-810 spectropolarimeter (Jasco). Spectra were recorded from 260 to 200 nm at 20°C in 1 mm path-length quartz cuvette at a protein concentration of 10 μM in 50 mM NaPO4 buffer at pH 7. Each CD spectrum was obtained by averaging 4 scans collected at a scan rate of 200 nm/min. Baseline spectra obtained with buffer were subtracted for all spectra.

### Preparation of liposomes and QCM-D measurements

DOPC and DOPG lipid mixtures were prepared in chloroform as described above except that desiccation was performed overnight. The lipids were rehydrated in 10 mM Tris pH 7.0, 100 mM NaCl, 5 mM MgCl_2_ buffer at a final concentration of 5 mg/mL using three consecutive cycles of freezing in liquid nitrogen and thawing in an ultrasonic bath (Merck). The rehydrated lipid solutions were extruded 21 times through a 100 nm diameter pore size polycarbonate membrane (Avanti Polar Lipids Inc.). The extruded solutions were stored at 4°C and consumed within a week after preparation.

A QCM-D E4 (QSense AB, Biolin Scientific AB, Gothenburg, Sweden) was used to measure MreB binding to planar supported lipid bilayers (SLBs) as previously reported for MinD and MinE (Renner & Weibel, 2012). Briefly, during QCM-D measurements, frequency and dissipation changes are recorded based on the piezoelectric properties of the crystal probe (Rodahl *et al*, 1995). The quartz crystals (QSense AB, Biolin Scientific AB, Gothenburg, Sweden) were coated with a custom 50 nm-thick layer of silicon dioxide by chemical vapor deposition (GeSiM GmbH, Dresden, Germany). Prior to each measurement, quartz crystals were thoroughly cleaned in a 1:1:5 volumetric ratio of concentrated ammonium hydroxide (Sigma-Aldrich), 30% hydrogen peroxide (Sigma Aldrich), and ultrapure water (Merck) at 70°C for 3 min. Prior to liposome deposition, the quartz crystals were then placed and oxidized in a plasma cleaner (Harrick Plasma, Ithaca, NY) for 2 min at high radio frequency. The oxidized (activated) crystals were placed into the QCM-D measurement chambers and immediately covered with 10 mM Tris buffer, 100 mM NaCl and 5 mM MgCl_2_. Subsequently, after a stable baseline was established, a liposome working solution (0.2 mg/mL) was pumped into the measurement chambers at 200 μL/min. After 2-20 min of incubation, a characteristic profile of supported planar lipid bilayer formation was observed (Fig. S7A) (Keller *et al*., 2000). After 5 min, the SLBs were rinsed with 10 mM Tris buffer containing 100 mM NaCl and 5 mM MgCl_2_ to remove unbound vesicles at 100 μL/min. The buffer was next exchanged to the reaction buffer (20 mM Tris-HCl pH 7, 500 mM KCl, 1 mM DTT and 2 mM EDTA). After a stable baseline was observed, MreB (±ATP, ADP, AMP-PNP) in reaction buffer (20 mM Tris-HCl pH 7, 500 mM KCl, 1 mM DTT, 5mM MgCl_2_) was added at 0.1, 0.5, 1 and 5 μM (low to high concentration) to the SLB at a pump speed of 100 μL/min for 5 min. The adsorption of MreB wild-type and mutants was measured for at least 20 min before exchanging and rinsing with reaction buffer for 5 min at 100 μL/min. In a series of experiments (from low to high MreB concentration), MreB was almost completely displaced by the rinsing step, allowing multiple adsorption steps on a single SLB. However, at higher MreB concentrations the rinsing was only partially effective (Fig. S7B). To avoid history effects on a SLB, we also reversed the MreB concentration steps (from high to low concentration). We calculated the thickness from frequency shifts using the Sauerbrey model included in the commercial analysis software tool QTools (QSense AB, Biolin Scientific AB, Gothenburg, Sweden). Each measurement was repeated at least twice with 2-3 repeats.

### NTPase activity assay

ATPase and GTPase activity of MreB were assayed by measuring the release of free inorganic phosphate (Pi) in a colorimetric assay using malachite green (Kodama *et al*, 1986; Mao *et al*, 2017). P_i_ produced was measured after a fix (end-point) or various (kinetic) incubation times in the reaction buffer (20 mM Tris, 500 mM KCl, 5 mM MgCl_2_) with appropriate supplements (e.g. 0.5 mM ATP or GTP, 0.05 mg/mL liposomes). The liposome solution was made on the day of the experiment by resuspending desiccated TE in water to 1 mg/mL. The reaction was initiated by the addition of MreB to the reaction mixture and ended by addition of 1 reaction volume of malachite revelation buffer (0.2 % (w/v) ammonium molybdate, 0.7 M HCl, 0.03 % (w/v) malachite green, 0.05 % (v/v) Triton X-100). Incubations were performed at 53°C and 37°C for 1h (end point) or less (kinetics). The quantity of P_i_ produced was determined by measuring the absorbance at 650 nm on a 96-well plate spectrophotometer (Synergy 2, Biotek). A mock reaction devoid of protein constituted the blank. A standard curve was made with a range of KH_2_PO_4_ diluted in the reaction buffer.

### Crystallization, structure determination and refinement

Freshly purified MreB^Gs^ containing an N-terminal His6-tag (stored in 20 mM Tris pH7, 500 mM KCl, 2 mM EDTA, 1 mM DTT) was concentrated by centrifugation using a vivaspin 5 000 MWCO membrane tube. All crystallization assays were performed at 293 K by sitting-drop vapor diffusion using facilities from the crystallization platform of I2BC. Crystals of apo MreB^Gs^ were obtained from a 100:100 nL mixture of protein at 3 mg/mL with a crystallization solution composed of 33% polyethylene glycol (PEG) 300 in 0.1 M MES pH 6.7. For co-crystallization assays, 10 mM ATP-Mg was added to 6 mg/mL of protein. Crystals of the complex were obtained with a crystallization solution containing 16% PEG 8000, 20% Glycerol and 0.04 M potassium phosphate. All crystals were flash-frozen in liquid nitrogen before data collection. Diffraction-quality crystals attained their full sizes in roughly 10-14 days.

Diffraction data were recorded on beam line PROXIMA 1 (synchrotron SOLEIL, France) at a wavelength of 0.9786 Å. Data were processed with the XDS package (Kabsch, 2010). All structures were solved by molecular replacement using the MOLREP program (Vagin & Teplyakov, 1997) using the crystal structure of MreB^Cc^ (PDB ID 4CZJ) (van den Ent *et al*., 2014), and the models were refined using PHENIX (Liebschner *et al*, 2019). The models were further improved by iterative cycles of manual rebuilding using COOT (Emsley *et al*, 2010). Final structural models were deposited in the Protein Data Bank (PDB) (Berman *et al*, 2000). Statistics for all the data collections, refinement of the different structures and the PDB codes are summarized in Table S1. All structural figures were generated with PyMOL (The PyMOL Molecular Graphics System, version 1.2r3pre, Schrödinger, LLC n.d.). Protein structure comparison was performed using the PDBeFold service at European Bioinformatics Institute (http://www.ebi.ac.uk/msd-srv/ssm) (Krissinel & Henrick, 2004). Protein interfaces, surfaces and assemblies were analyzed using the PDBePISA service at European Bioinformatics Institute (http://www.ebi.ac.uk/pdbe/prot_int/pistart.html) (Krissinel & Henrick, 2007).

## Acknowledgments

We thank Davy Martin for CD acquisitions, Human Rezai and Juan Hermoso for useful discussions and Xavier Henry for useful contributions upstream this work. This project has received funding from the European Research Council (ERC) under the European Union’s Seventh Framework Program (FP7) and the Horizon 2020 research and innovation program (grant agreement No 311231 and grant agreement No 772178, respectively, to R.C.-L.). We also thank the Labex Cell(n)Scale (ANR-11-LABX0038), Paris Sciences et Lettres (ANR-10-IDEX-0001-02), and the Cell and Tissue Imaging (PICT-IBiSA), Institut Curie, member of the French National Research Infrastructure France-BioImaging (ANR-10-INBS-04). L.D.R. acknowledges funding by the VolkswagenStiftung. This work benefited from the expertise of Christine Péchoux at the MIMA2 facility (Université Paris-Saclay, INRAE, AgroParisTech, GABI, 78350, Jouy-en-Josas, France) for TEM observations https://doi.org/10.15454/1.5572348210007727E12, and of the crystallization platform of I2BC, supported by the French Infrastructure for Integrated Structural Biology (FRISBI, ANR-10-INSB-05-05). We acknowledge the synchrotrons ESRF (Grenoble, France) and SOLEIL (Saint-Aubin, France) for provision of synchrotron radiation facilities and we would like to thank the staffs of beamlines PROXIMA-2A and PROXIMA-1 at SOLEIL, and 1D23-2 at ESRF for assistance and advices during data collection.

## Authors contributions

**Wei Mao:** Formal analysis; investigation; visualization; writing – original draft. **Lars D. Renner:** Formal analysis; investigation; methodology; validation; visualization; writing – review & editing. **Charlène Cornilleau:** Investigation. **Ines Li de la Sierra-Gallay:** Formal analysis; investigation, visualisation. **Sarah Benlamara:** Investigation. **Yoan Ah-Seng:** Investigation. **Herman Van Tilbeurgh:** Methodology. **Sylvie Nessler:** Data curation; formal analysis; supervision; validation; visualization; writing – review & editing. **Aurélie Bertin:** Formal analysis; investigation; methodology; validation; visualization; writing – review & editing. **Arnaud Chastanet:** Conceptualization; data curation; formal analysis; investigation; methodology; supervision; validation; visualization; writing – original draft; writing – review & editing. **Rut Carballido-López:** Conceptualization; formal analysis; funding acquisition; project administration; supervision; validation; writing – original draft; writing – review & editing.

In addition to the [CRediT](https://credit.niso.org/) individual author contributions listed above, the general contributions are:

R.C.-L. administrated and acquired the financial support for the project leading to this publication; A.C and R.C.-L. conceptualized, designed the work and supervised the project; L.D.R., I. L.d.l.S.-G., S.N., A.B, A.C. and R.C.-L. conceived experiments and validated results and experiments; W.M., L.D.R., C.C., I.d-G., S.B., A.B. and A.C., performed experiments or data collection; Y.A.-S. performed evidence collection; W.M., L.D.R., I. L.d.l.S.-G., S.N., A.B., A.C. and R.C.-L. analyzed and/or interpreted data; H.V.-T. contributed to scientific discussions; A.C. and R.C.-L. wrote a complete original draft and revised and edited the manuscript; W.M. contributed to the writing of part of an original draft; L.D.R., S.N. and A.B. contributed to the writing and review of the manuscript. All authors approved the manuscript and its contents.

## Supporting Information

Supplementary Figures S1 to S12

Supplementary Tables S1 to S5

## Supplementary Figure Legends

**Figure S1.**
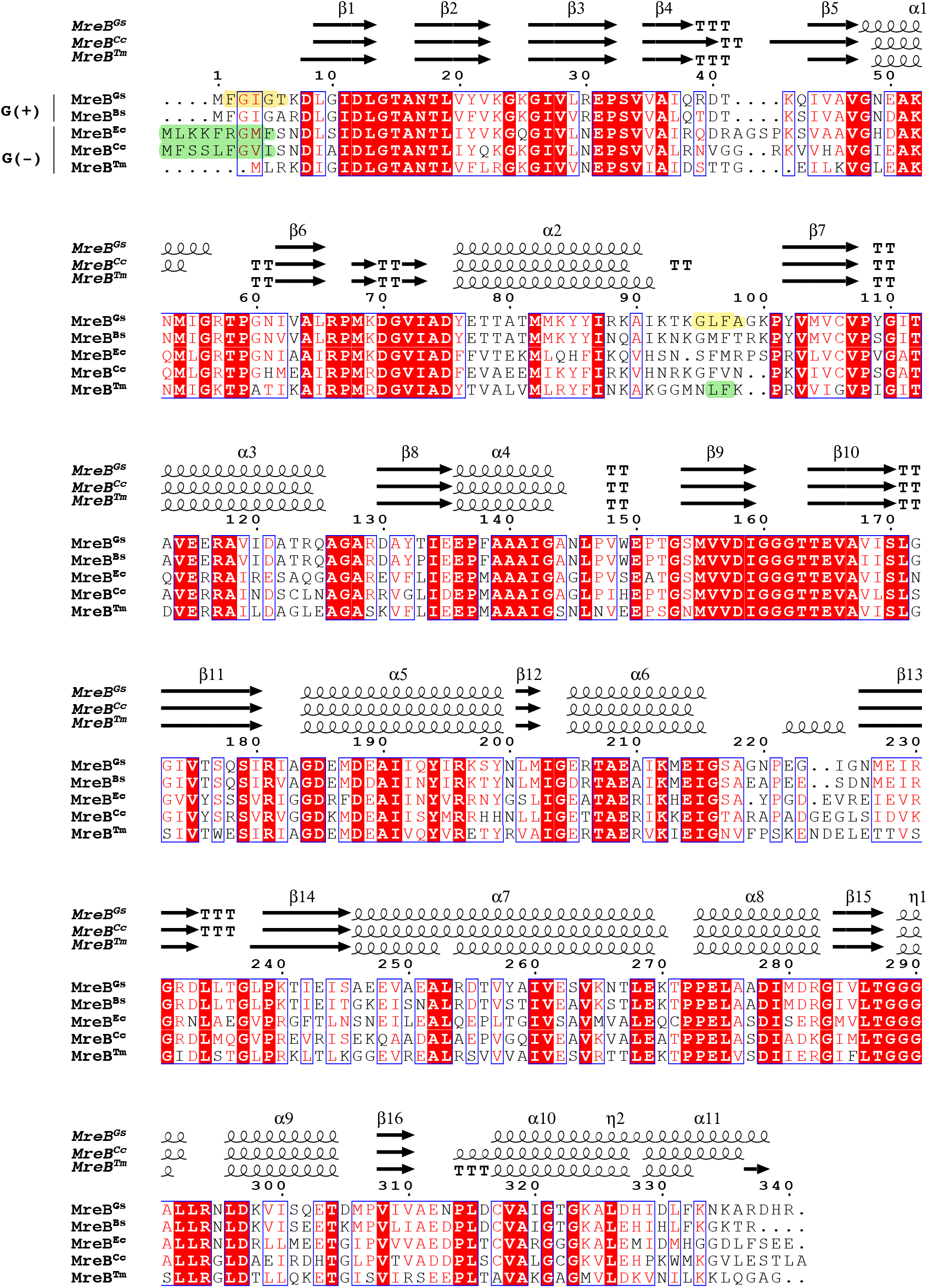
Multiple sequence alignment of MreB proteins. The sequence of from *G. stearothermophilus* (MreB^Gs^) was aligned using Clustal-Ω at PRABI (https://npsa-prabi.ibcp.fr/cgi-bin/npsa_automat.pl?page=/NPSA/npsa_clustalw.html) against the homologous MreB sequences of the G+ bacterium *B. subtilis* (MreB^Bs^, GenBank ID ATA60829.1) and the G- bacteria *E. coli* (MreB^Ec^, GenBank ID P_417717), *C. crescentus* (MreB^Cc^, GenBank ID YP_002516985.1) and *T. maritima* (MreB^Tm^, GenBank ID AAD35673.1), respectively. Sequence numbering is relative to MreB^Gs^. Secondary structure information extracted from the crystal structures of MreB^Gs^ (PDB ID 7ZPT), MreB^Cc^ (PDB ID 4CZM) and MreB^Tm^ (PDB ID IJCE) are indicated above the sequences using ESPript (Robert & Gouet, 2014). Beta strands are numbered β1 to β16, alpha helices α1 to α11, and 3_10_ helices η1 to η2, according to the MreB^Gs^ structure.α- and β-turns are depicted as TTT and TT, respectively. Blue frames indicate homologous regions, similar residues are indicated in red and identical residues in white on red background. The residues of the amino-terminus and α2β7 hydrophobic loops deleted in the mutants ΔNter and ΔGLFA of MreB^Gs^ are highlighted in yellow. Amphipathic helices in MreB^Ec^ and MreB^Cc^ and the hydrophobic loop in MreB^Tm^ are highlighted in green.

**Figure S2.**
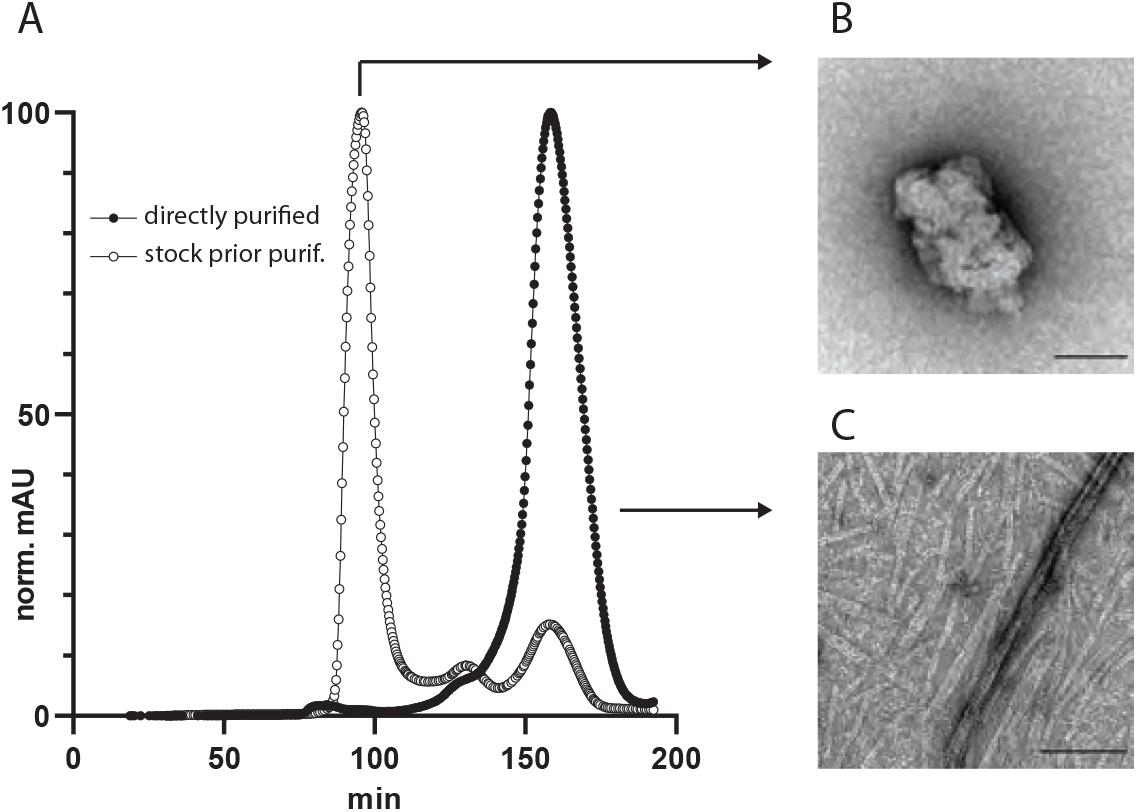
(**A**) Typical size exclusion chromatography elution profiles of MreB^Gs^. MreB^Gst^ (wild-type) was loaded on a HiLoad™ 16/600 Superdex™ 200 pg (GE healthcare) size exclusion column immediately after elution from a Nickel-NTA affinity purification column (plein circles) or after a subsequent 4°C overnight incubation (empty circles). When size exclusion chromatography was performed using freshly purified protein from the affinity column, MreB^Gs^ (37.36 kDa) eluted mainly as a single peak corresponding to the monomeric form of the protein according to the calibration of the column. In contrast, overnight conservation of the eluate before loading onto the Superdex column leads to the irreversible formation of high molecular weight assemblies (aggregates) eluting at the dead volume of the column. (**B, C**) TEM micrographs of negatively stained samples of MreB^Gs^ from the elution fractions indicated with an arrow. High molecular weight (=short retention time; B) or monomeric (=long retention time; C) MreB^Gs^ forms were incubated in conditions supporting polymerization and mounted on EM grids for TEM observation. High molecular weight MreB^Gs^ forms are aggregates of various sizes and shapes independent on the conditions tested (B). Monomeric MreB^Gs^ polymerizes into pairs and sheets of protofilaments (C).

**Figure S3.**
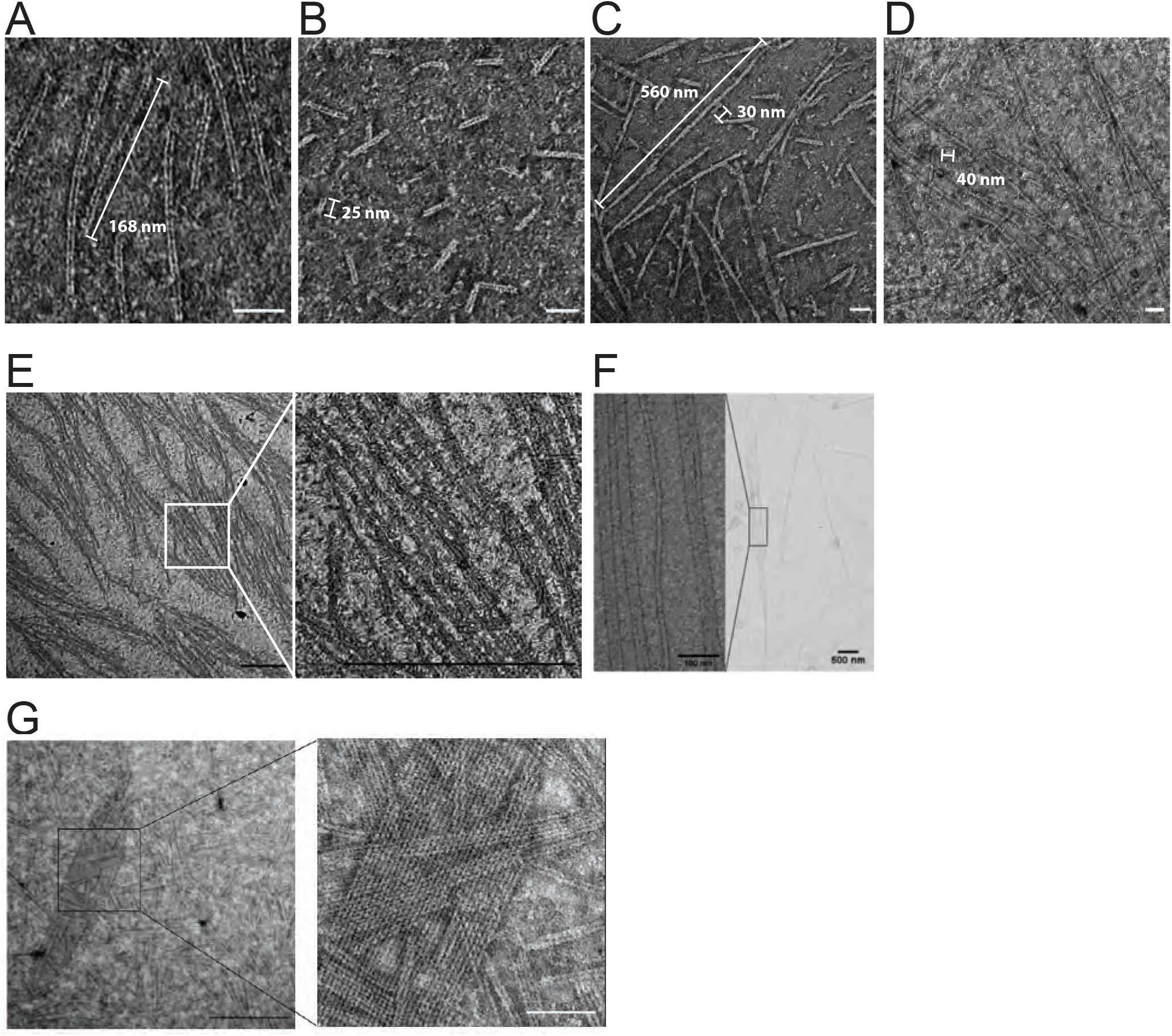
MreB^Gs^ polymers display a broad range of lengths. **(A-D)** Dual protofilaments of MreB^Gs^ observed on various fields of a single EM grid. Example of fields containing exclusively medium size polymers (> 100 nm) (A); exclusively short polymers (< 50 nm) (B); a mix of medium (some bundling) and short polymers (C), and a mix of long (> 1 μm) and short polymers (D). In D, the long polymers are extending beyond the edges of the field of view. Scale bars, 50 nm. **(E, F)** MreB^Gs^ polymers can form filaments extending over several μm. Scale bars, 500 nm. **(G)**MreB^Gs^ polymers can associate laterally to form sheets of various widths. Scale bars, 500 nm (black) and 100 nm (white).

**Figure S4.**
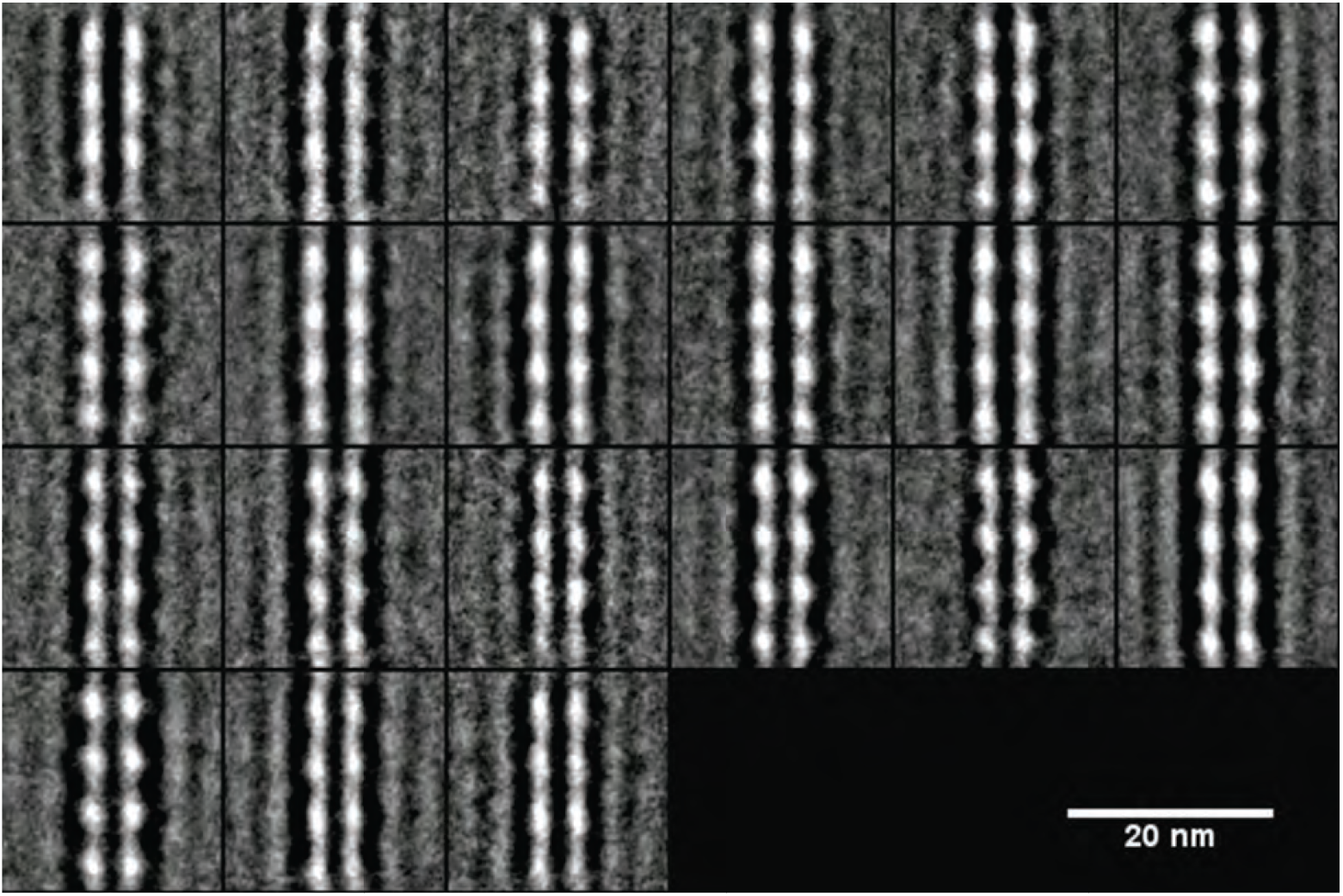
2D averaging of negatively stained images of MreB^Gs^ dual protofilaments showing the symmetrical arrangement of monomers. Displayed are the 21 classes of images generated by 2D image processing (alignment and classification from 1 554 individual raw image). Scale bar, 20 nm.

**Figure S5.**
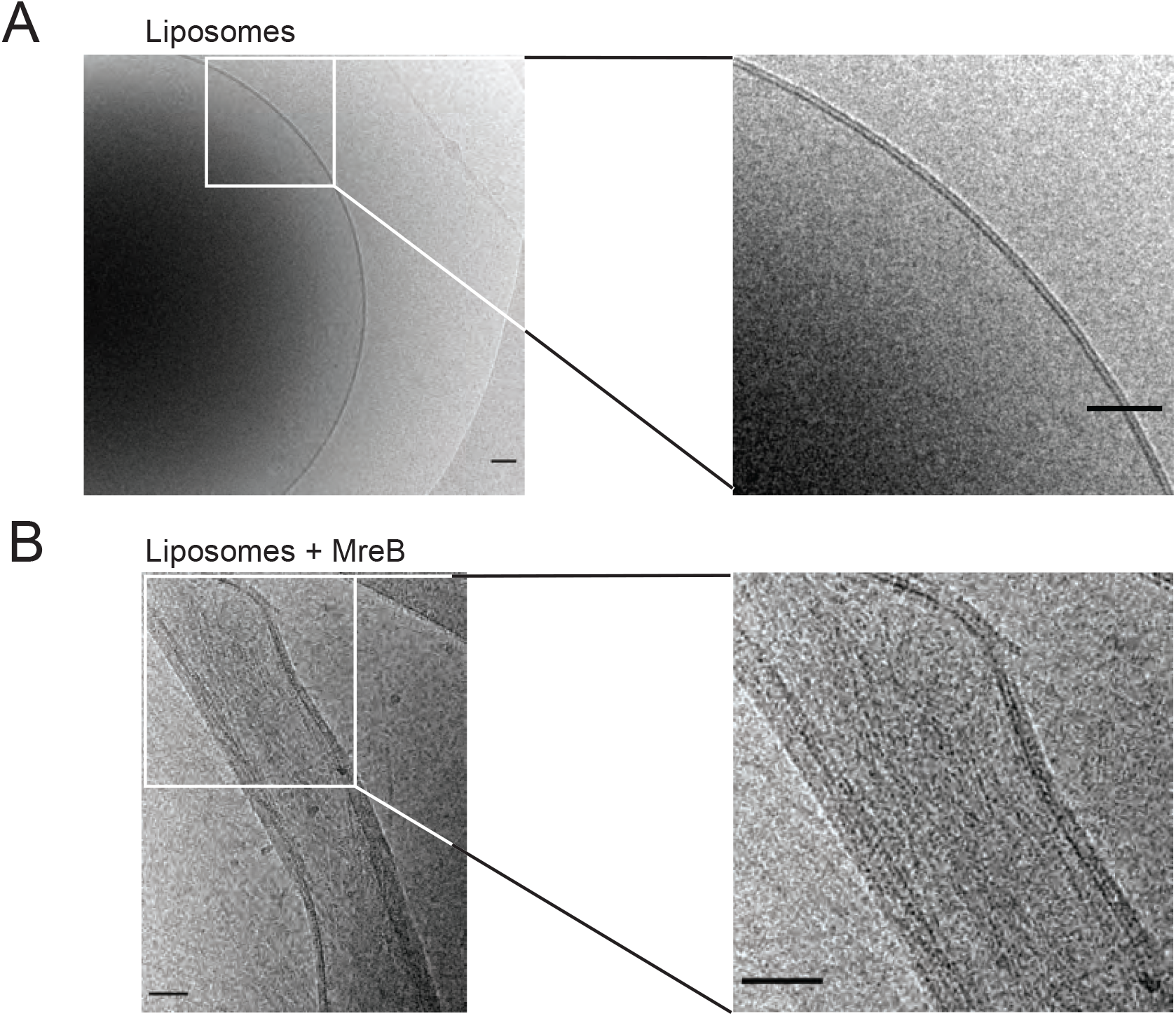
MreB^Gs^ polymers coat and distort liposomes. Cryo-EM micrographs of 0.37 mg/mL liposomes made from lipid total extract from *E. coli* alone, shown as negative control (**A**), or mixed with 0,05 mg/mL purified MreB^Gs^ in the presence of 2mM ATP and incubated for 2h at room temperature (**B**). MreB^Gs^ extensively coated the liposomes and deformed them into faceted vesicles. Scale bars, 50 nm.

**Figure S6.**
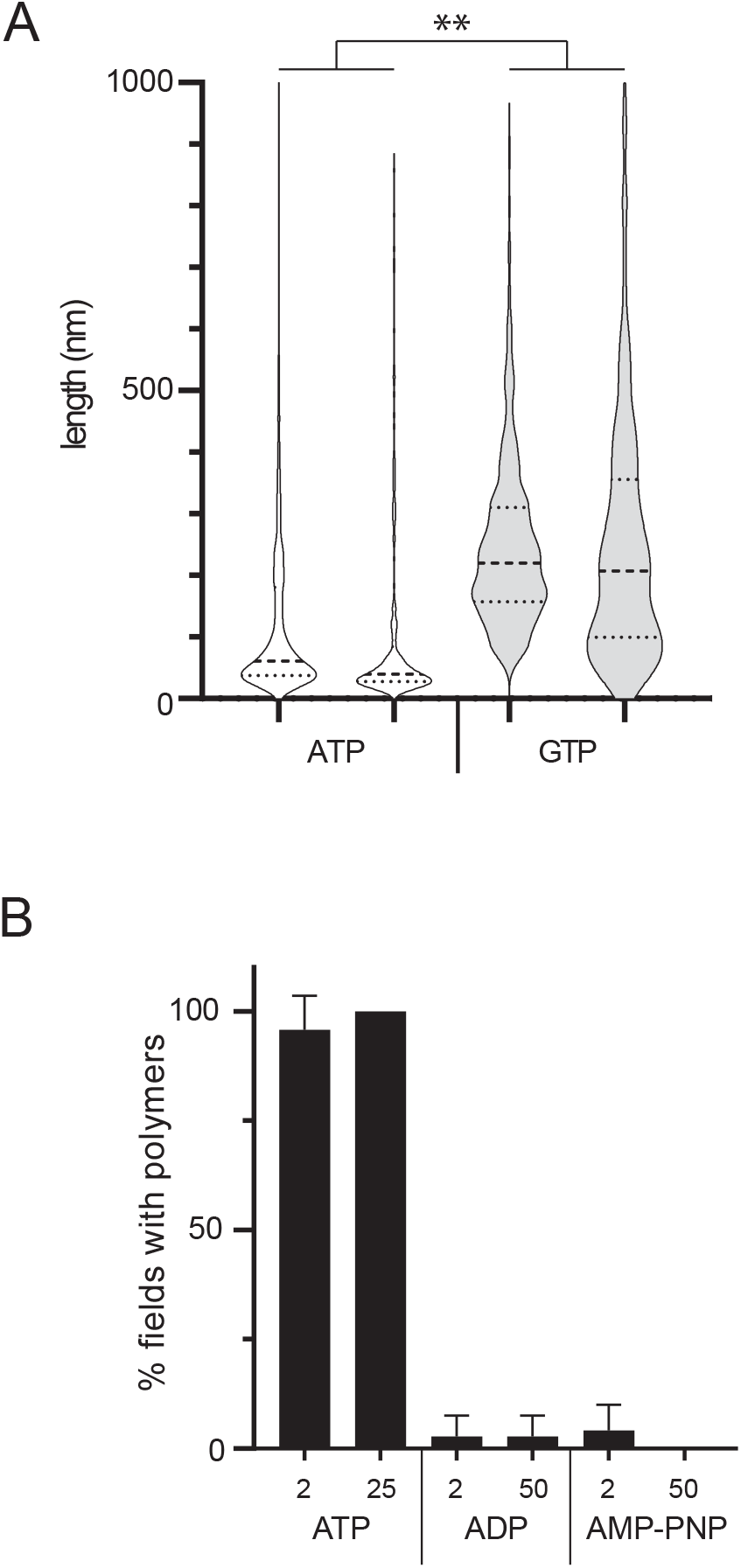
(**A**) Size distribution of MreB^Gs^ double filaments set to polymerize in the presence of ATP or GTP (2mM) in otherwise standard polymerization conditions. Negative stained EM micrographs were analyzed under FIJI and the length of filaments < 1 μm were individually measured. Values are distributions of length of at least 800 filaments per condition from 2 independent experiments. Median (dashed lines) and quartiles (dotted line) are displayed. The difference between the two conditions are significantly different in a nested T-test (P-value = 0.006). (**B**) Quantification of MreB^Gs^ polymer formation in the presence of high concentrations of nucleotides. MreB^Gs^ was set to polymerize in the presence of ATP (2 and 25 mM), ADP (2 and 50 mM) or AMP-PNP (2 and 50 mM) in otherwise standard polymerization conditions. EM images were acquired on 12 randomly picked position per EM grid, spread over the entire grids. Images were categorized based on the sole presence or absence of polymers. Values are average of at least two independent experiments. Error bars are standard deviations.

**Figure S7.**
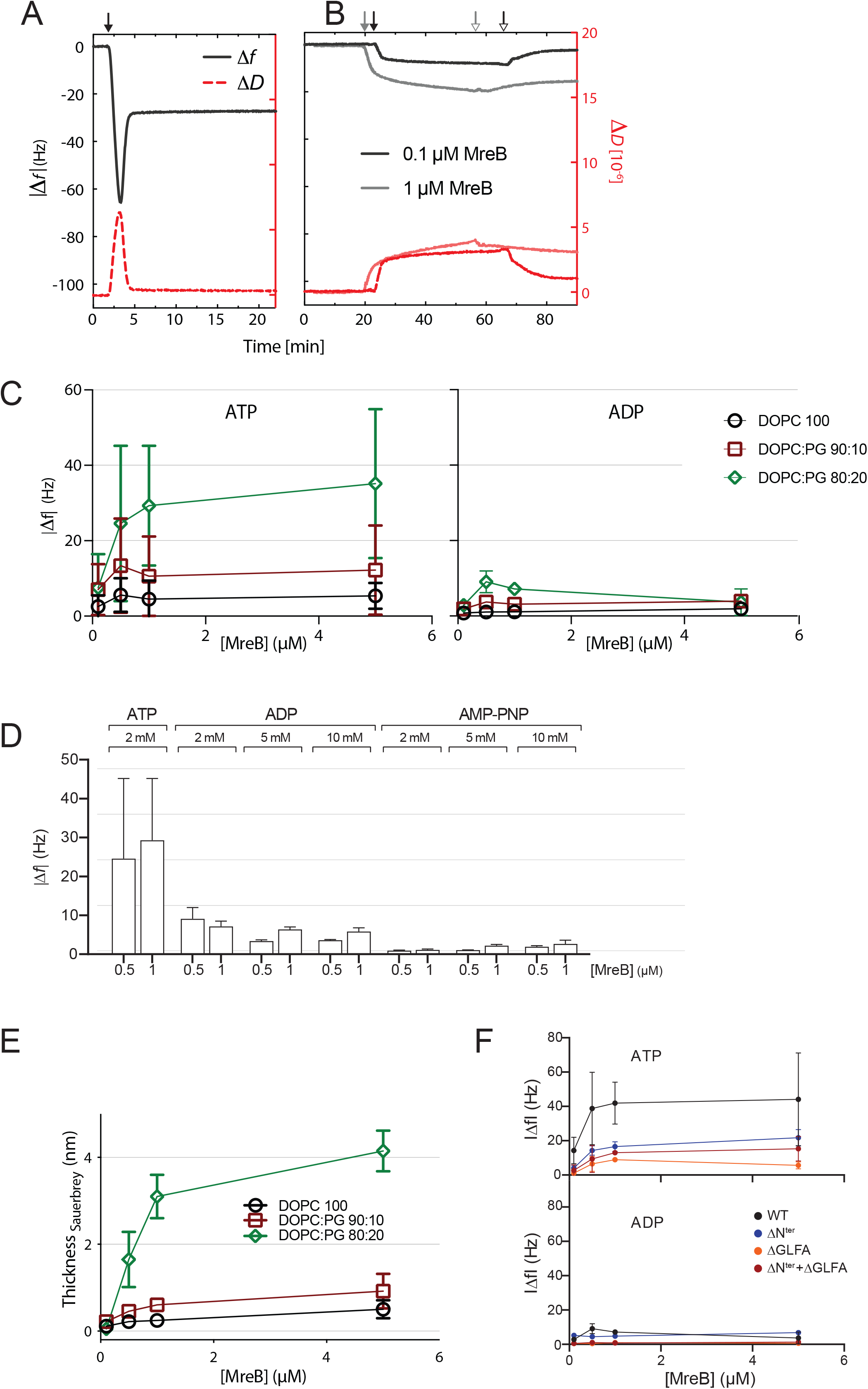
QCM-D experiments of MreB^Gs^ adsorption on supported lipid bilayers. **(A)** Lipid bilayer formation on crystal with SiO_2_ layers. Supported lipid bilayers (SLBs) are formed by spontaneous rupture of adsorbed liposomes as indicated by frequency shifts (Δ*f*, black solid lines) and dissipation shifts (Δ*D*, red dotted lines). Exemplarily shown is the formation of DOPC:DOPG 90:10 SLB from DOPC:DOPG 90:10 liposomes. The solid black arrow indicates the addition of liposomes to the SiO_2_ surface. (**B**) Subsequently, MreB^Gs^ wild-type in various concentrations (here 0.1 μM, black line and 1 μM, grey line) are added to SLBs. The closed and open arrows indicate the start and end of the protein addition (followed by rinsing with polymerization buffer), respectively. **(C)** MreB^Gs^ binds to SLBs of different lipid ratios in the presence of ATP but not in the presence of ADP. Error bars are standard deviations of n=3. **(D)** High concentrations of ADP and AMP-PNP do not support adsorption of MreB^Gs^ to SLBs. Frequency changes were measured for the binding of purified MreB^Gs^ to SLBs in QCM-D experiments. Incubations were performed in polymerization buffer containing 2mM ATP or the indicated concentrations of ADP or AMP-PNP. SLBs consisted of DOPC:DOPG 80:20. Values are averages of at least two independent experiments. Error bars represent standard deviations of n≥2. **(E)** Thickness of the MreB protein layer on the SLBs calculated with the Sauerbrey equation. **(F)** The hydrophobic α2-β7 loop and the N-terminus domain of MreB^Gs^ enhance adsorption to the SLB, in an ATP-dependent manner. Frequency change (|Δ*f*|) measured for the binding of various concentrations (0.1 - 5 μM) of purified wild-type (WT) and mutant forms of MreB^Gs^ to SLBs, assayed by QCM-D. Incubations were performed in polymerization buffer containing 2 mM ATP or ADP. SLBs contained an 80:20 molecular ratio of DOPC:DOPG. Error bars are standard deviations of n=3.

**Figure S8.**
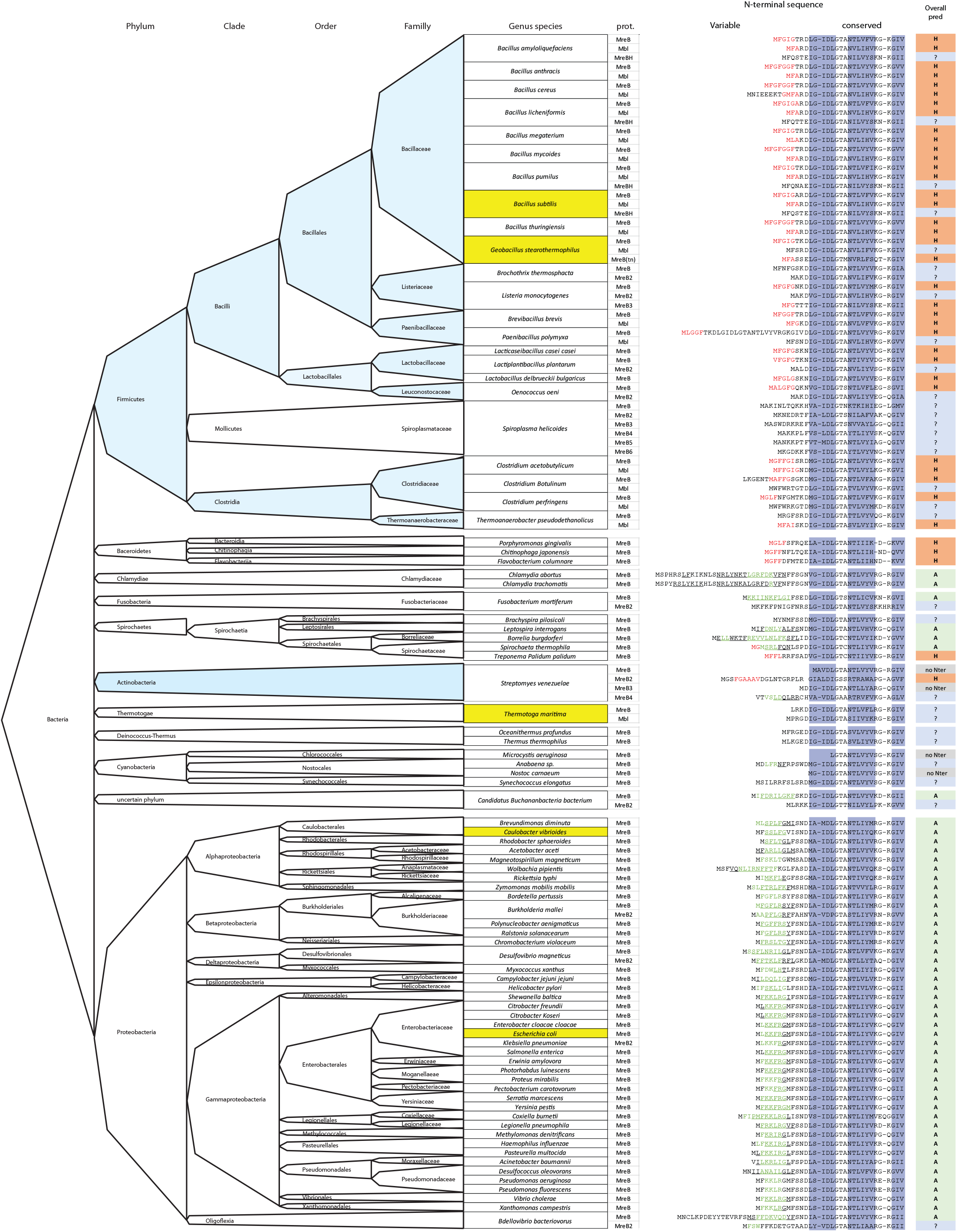
Distribution of N-terminal amphipathic helix and hydrophobic sequences in the bacterial kingdom. N-terminal sequences of MreB proteins from selected species across the bacterial kingdom were aligned using Clustal-Ω. The N-terminal sequences were analyzed for the presence of putative α-helix (underscore) and/or amphipaticity (green) using the Amphipaseek tool at Prabi (https://npsa-prabi.ibcp.fr/), and for the presence of hydrophobic sequences (red). Dark blue columns mark the β-sheets 1, 2 and 3 according to MreB^Gs^ structure (Fig. S1). The prediction for putative anchoring structures is summarized in the right column: A (green), amphipathic helix; H (red), hydrophobic sequence; ? (blue), unknown. Species of interest aligned in Fig. S1 are highlighted in yellow. G+ bacteria (with low and high GC %) are colored in light blue.

**Figure S9.**
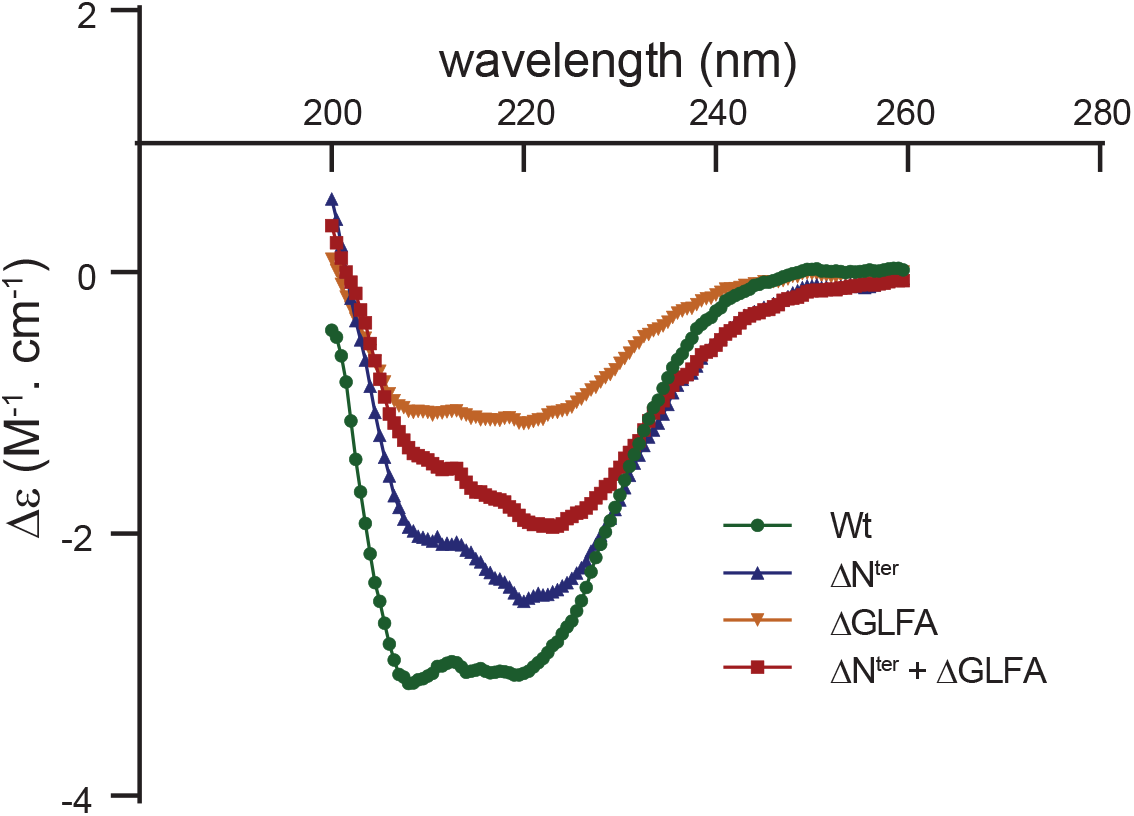
Circular dichroism (CD) spectra showing similar folding of the wild-type and the deletion mutants of the α2-β7 (ΔGLFA), the N-terminus (ΔN^ter^) or both domains (ΔN^ter^+ΔGLFA) of recombinant MreB^Gs^.

**Figure S10.**
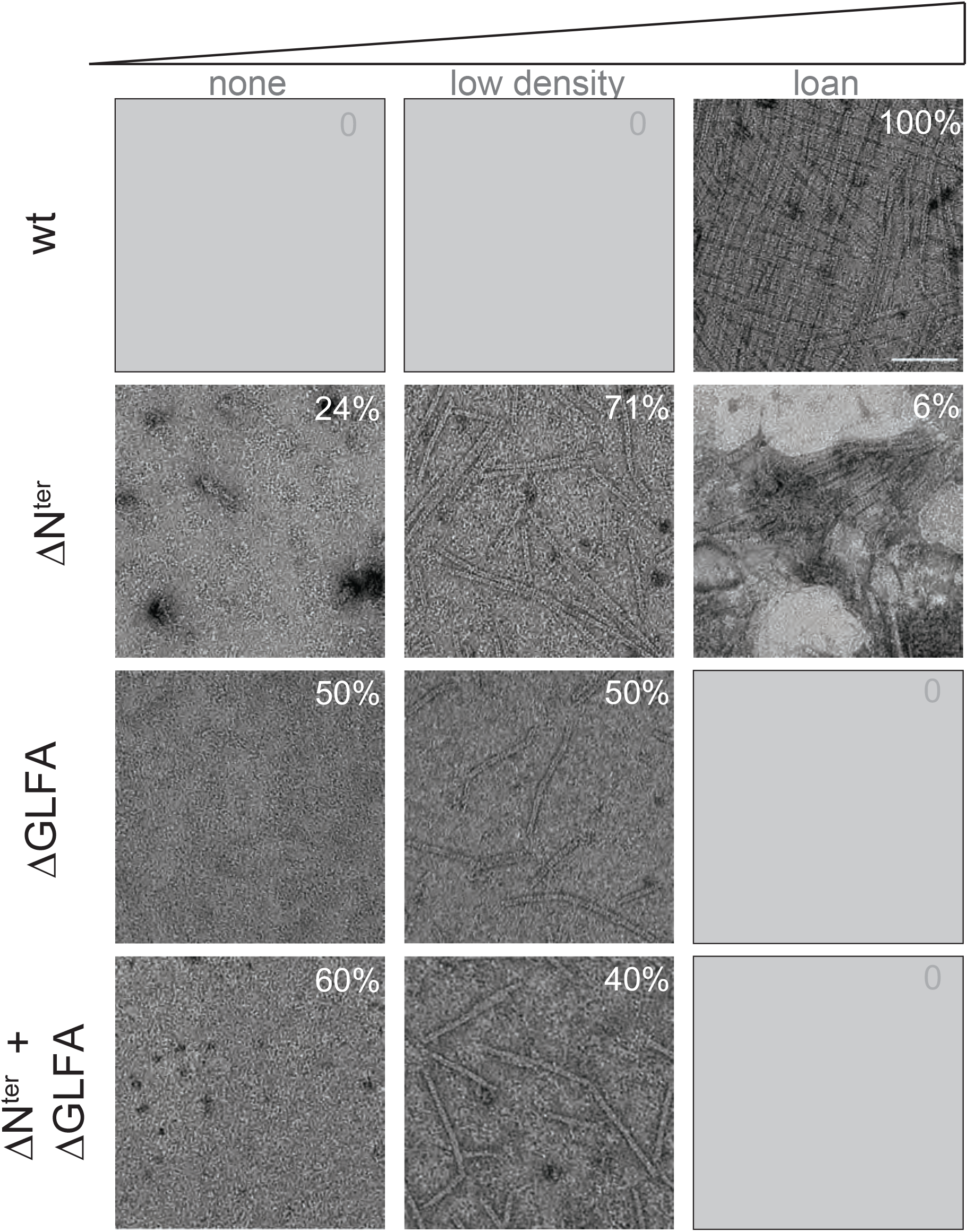
Deletion of the amino-terminal sequence, the GLFA residues of the α2-β7 hydrophobic loop or both, decrease the quantity of MreB^Gs^ polymers on a lipid monolayer. Because the repartition of the polymers on TEM grids are heterogeneous, we acquired for each of two experimental replicas, images on 12 random locations widespread on the entire grids. Images were subsequently distributed based on the presence of polymers: none, low density, or loan. Here are presented zoomed-in regions of the grids with typical examples of each category of images for the wild type MreB^Gs^ protein (wt) and the three deletion mutants (deleted for the amino-terminus (ΔN^ter^), for the hydrophobic loop (ΔGLFA) or for both (ΔN^ter^+ΔGLFA)). Grey panels indicate that no images were found for that category. Numbers indicate the percentage of observed images (sum of replicates) of each category for each protein. Scale bar, 100 nm.

**Figure S11.**
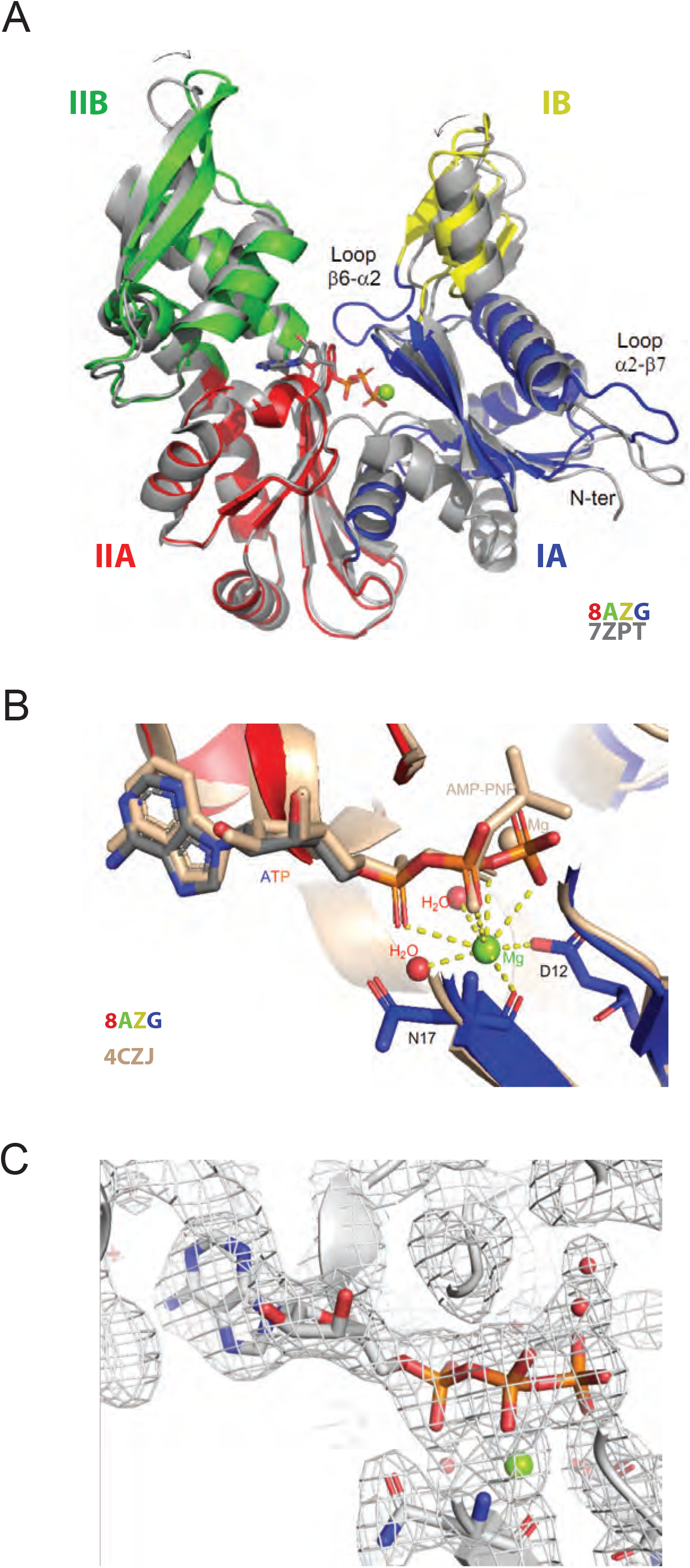
Crystal structure of MreB^Gs^ bound to ATP-Mg. (**A**) Comparison of the ATP-Mg-bound form with the apo form of MreB^Gs^. One subunit of ATP-bound MreB^Gs^ (PDB ID 8AZG), colored by subdomains, is superimposed with the apo form of the protein (PDB ID 7ZPT), colored in gray. The bound nucleotide is shown as sticks colored by atom type (C in grey, N in blue, O in red and P in orange). The associated magnesium ion is shown as a green sphere. Loop β6-α2, stabilized by the presence of the bound nucleotide, is labeled, as well as loop α2-β7 and the N-terminus, which display alternative conformations. (**B**) Comparison of the ATP-Mg-bound MreB^Gs^ with MreB^Cc^ bound to AMP-PNP/Mg. Close view of the superimposed nucleotide-binding sites. MreB^Gs^ (PDB ID 8AZG) and MreB^Cc^ (PDB ID 4CZJ) are shown as cartoon colored by domain and in beige, respectively. The nucleotide molecules are shown as sticks. The bound ATP/Mg is colored by atom type (C in gray, N in blue, O in red, P in orange and Mg in green). The bound AMP-PNP/Mg is colored in beige. Two conserved residues (D12 and N17) involved in the coordination of the Mg^2+^ ion in the MreB^Gs^ complex are shown as sticks and labeled. Two water molecules also involved in Mg^2+^ coordination are shown as red spheres. (**C**) Electron density map of ATP-Mg bound to MreB^Gs^. The 2Fo-Fc map contoured at 1.2 σ of the nucleotide binding site is shown as a gray mesh.

**Figure S12.**
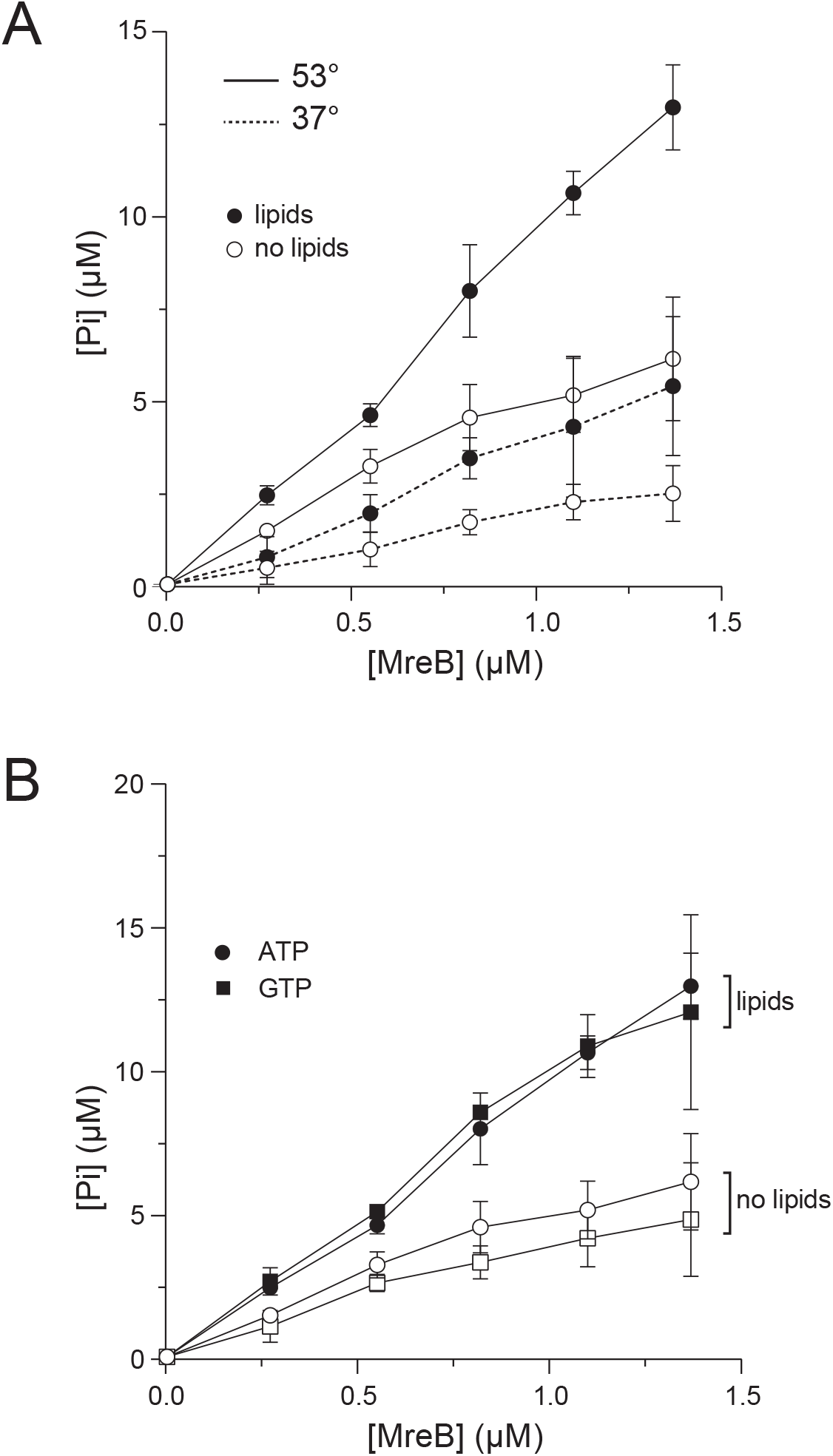
(**A**) The ATPase activity of MreB^Gs^ is stimulated at high temperature. Release of P_i_ detected by malachite green assay for a range of MreB^Gs^ concentrations (0.27 – 1.37 μM) in the presence or absence of 0.05 mg/mL liposomes in polymerization buffer (0.5 mM ATP) after 1 h incubation at 53°C or 37°C. Error bars are standard deviations of at least two independent measurements. (**B**) MreB shows a similar hydrolytic activity toward GTP and ATP and is stimulated in the presence of lipids. Release of P_i_ detected by malachite green assay in the presence of ATP or GTP (0.5 mM), after 1 h incubation at 53°C in the presence or absence of 0.05 mg/mL liposomes for a range of MreB^Gs^ concentrations (0.27 – 1.37 μM). Error bars are standard deviations of at least two independent measurements.

**Table S1.**
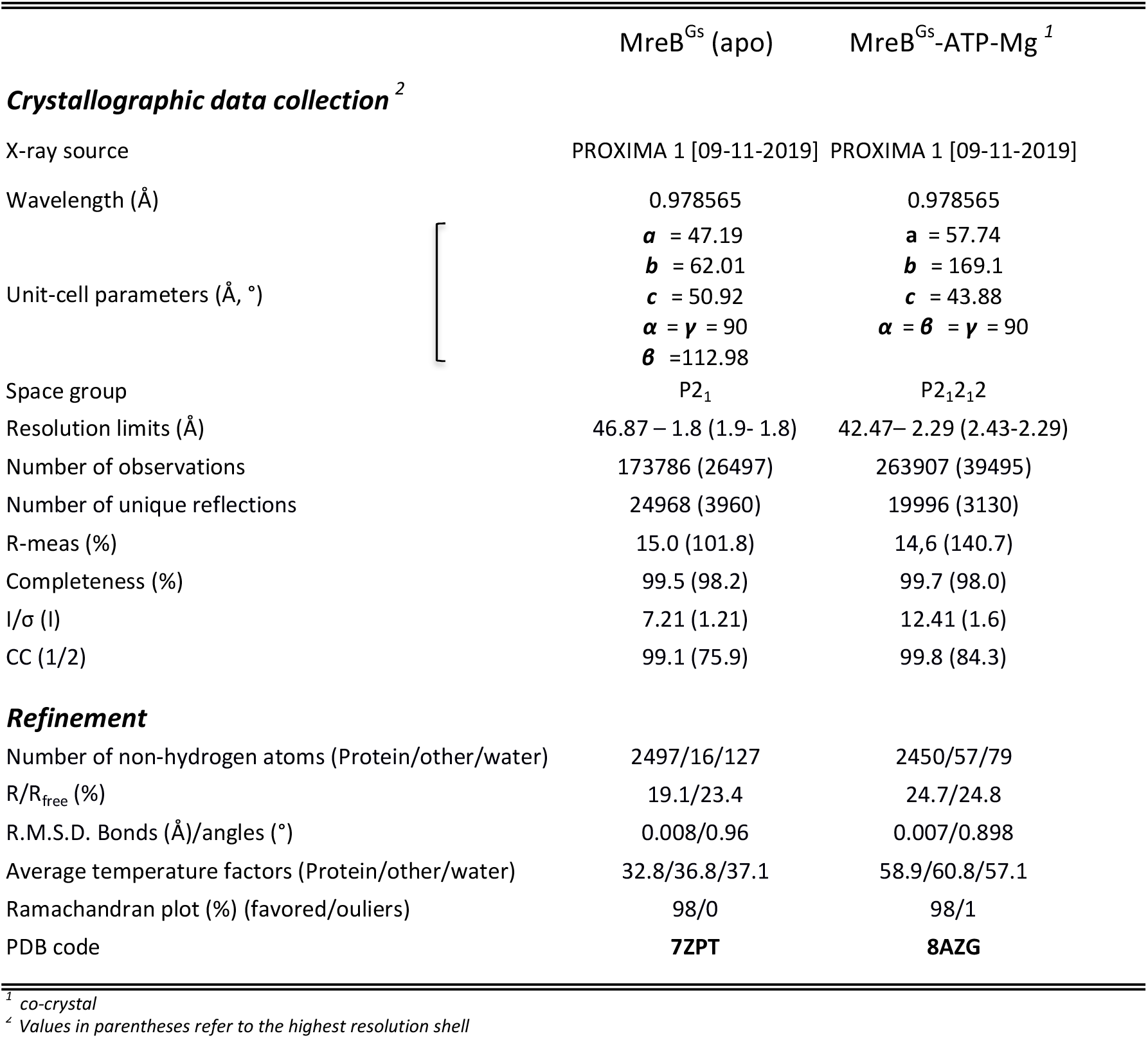
Data-collection and refinement statistics

**Table S2.**
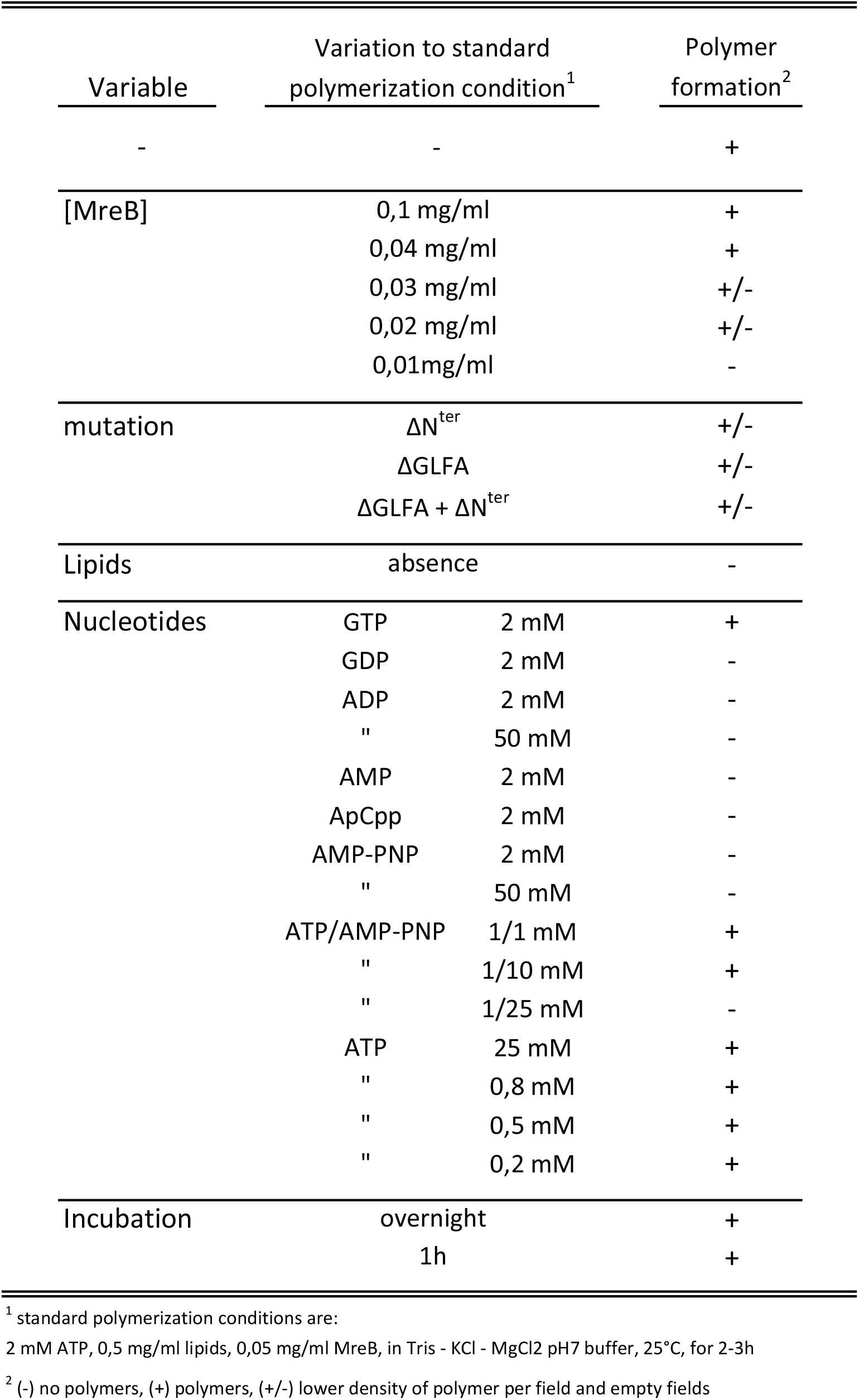
List of polymerization condition assayed

| Variable | Variation to standard polymerization condition<sup>1</sup> |  | Polymer formation<sup>2</sup> |
| --- | --- | --- | --- |
| - | - |  | + |
| [MreB] | 0,1 mg/ml |  | + |
|  | 0,04 mg/ml |  | + |
|  | 0,03 mg/ml |  | +/- |
|  | 0,02 mg/ml |  | +/- |
|  | 0,01mg/ml |  | - |
| mutation | $\Delta N^{\text{ter}}$ | | +/- |
| | $\Delta \text{GLFA}$ | | +/- |
| | $\Delta \text{GLFA} + \Delta N^{\text{ter}}$ | | +/- |
| Lipids | absence |  | - |
| Nucleotides | GTP | 2 mM | + |
|  | GDP | 2 mM | - |
|  | ADP | 2 mM | - |
|  | " | 50 mM | - |
|  | AMP | 2 mM | - |
|  | ApCpp | 2 mM | - |
|  | AMP-PNP | 2 mM | - |
|  | " | 50 mM | - |
|  | ATP/AMP-PNP | 1/1 mM | + |
|  | " | 1/10 mM | + |
|  | " | 1/25 mM | - |
|  | ATP | 25 mM | + |
|  | " | 0,8 mM | + |
|  | " | 0,5 mM | + |
|  | " | 0,2 mM | + |
| Incubation | overnight |  | + |
|  | 1h |  | + |

<sup>1</sup> standard polymerization conditions are:
2 mM ATP, 0,5 mg/ml lipids, 0,05 mg/ml MreB, in Tris - KCl - MgCl<sub>2</sub> pH7 buffer, 25°C, for 2-3h
<sup>2</sup> (-) no polymers, (+) polymers, (+/-) lower density of polymer per field and empty fields

**Table S3.** Proteins used in this study

| name | MreB<sup>Gs</sup> * | expression vector |
| --- | --- | --- |
| WT | MHHHHHH M<sup>1</sup>FGIGTKDLGI<sup>11</sup> (...) K<sup>94</sup>GLFAGK<sup>100</sup> (...) | pCC110 |
| ΔN<sup>ter</sup> | MHHHHHH M<sup>1</sup>DLGI<sup>11</sup> (...) K<sup>94</sup>GLFAGK<sup>100</sup> (...) | pCC116 |
| ΔGLFA | MHHHHHH M<sup>1</sup>FGIGTKDLGI<sup>11</sup> (...) K<sup>94</sup>GK<sup>100</sup> (...) | pCC117 |
| ΔN<sup>ter</sup> | MHHHHHH M<sup>1</sup>DLGI<sup>11</sup> (...) K<sup>94</sup>GK<sup>100</sup> (...) | pCC115 |

\*numbering of aminoacids refers to *MreB*<sup>Gs</sup> wt sequence

**Table S4.** Strains used in this study

| Strain | Relevant genotype | information | Source or reference<sup>1</sup> |
| --- | --- | --- | --- |
| <u>*G. stearothermophilus*</u> |  |  |  |
| ATCC 7953 | wild type isolate |  | BGSC (Ref. W9A12) |
| <u>*E. coli*</u> |  |  |  |
| T7 express | *F- λ- fhuA2 [lon] ompT lacZ ::T7 gene1 gal sulA11 Δ(mcrC-mrr)114 ::IS10 R (mcr-73 ::miniTn10-TetS)2 R(zgb-210 ::Tn10)(TetS) endA1 [dcm]* | expression strain carrying the T7 RNA polymerase gene into the *lac* operon, allowing controlled IPTG-induced expression of a gene of interest | New England Biolabs |
| EcRCL2 | pET28a | replicative plasmid pET28a(+) (Novagen) allowing IPTG-induction of a gene of interest; in DH10b | lab collection |
| EcRCL212 | pCC110::(*mreB*<sup>Gs</sup>) | pET28a(+) derivative carrying a wild type copy of *mreB*<sup>Gs</sup> under control of the T7 promoter | pCC110 → T7 express |
| EcRCL243 | pCC115::(*mreB*<sup>Gs</sup> δ 2->7;δGLFA) | pCC110 derivative carrying a copy of *mreB*<sup>Gs</sup> deleted for codons 2-7 (FGIGTK) and 102-105 (GLFA) | pCC115 → T7 express |
| EcRCL244 | pCC116::(*mreB*<sup>Gs</sup> δ 2->7) | pCC110 derivative carrying a copy of *mreB*<sup>Gs</sup> deleted for codons 2-7 (FGIGTK) | pCC116 → T7 express |
| EcRCL245 | pCC117::(*mreB*<sup>Gs</sup> δGLFA) | pCC110 derivative carrying a copy of *mreB*<sup>Gs</sup> deleted for codons 102-105 (GLFA) | pCC117 → T7 express |

1: Arrows indicate construction by transformation with chromosomal or plasmidic DNA

**Table S5.**
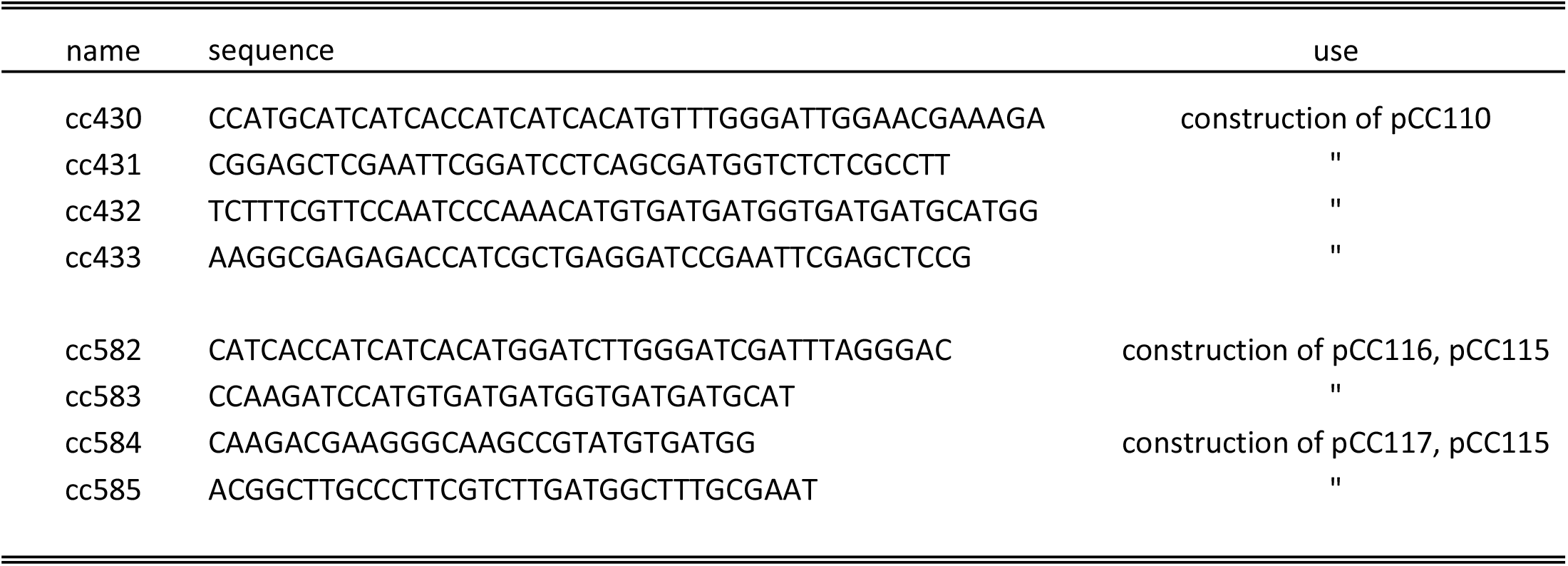
Oligonucleotides used in this study

